# Astrocyte-neuron crosstalk through Hedgehog signaling mediates cortical circuit assembly

**DOI:** 10.1101/2020.07.15.204263

**Authors:** Yajun Xie, Aaron T. Kuan, Wengang Wang, Zachary T. Herbert, Olivia Mosto, Olubusola Olukoya, Manal Adam, Steve Vu, Minsu Kim, Nicolás Gómez, Diana Tran, Claire Charpentier, Ingie Sorour, Michael Y. Tolstorukov, Bernardo L. Sabatini, Wei-Chung Allen Lee, Corey C. Harwell

## Abstract

Neuron-glia relationships play a critical role in the regulation of synapse formation and neuronal specification. The cellular and molecular mechanisms by which neurons and astrocytes communicate and coordinate are not well understood. Here we demonstrate that the canonical Sonic hedgehog (Shh) pathway is active in cortical astrocytes, where it acts to coordinate layer-specific synaptic connectivity and functional circuit development. We show that Ptch1 is a Shh receptor that is expressed by cortical astrocytes during development and that Shh signaling is necessary and sufficient to promote the expression of layer-specific astrocyte genes involved in regulating synapse formation and function. Loss of Shh in layer V neurons reduces astrocyte complexity and coverage by astrocytic processes in tripartite synapses, moreover, cell-autonomous activation of Shh signaling in astrocytes promotes cortical excitatory synapse formation. Together, these results suggest that Shh secreted from deep layer cortical neurons acts to specialize the molecular and functional features of astrocytes during development to shape circuit assembly and function.

## INTRODUCTION

The mammalian cerebral cortex is organized into six layers composed of neurons that share common projection patterns, molecular identities and intrinsic physiological properties (Custo Greig et al. 2013). Deep layer (layer V and VI) cortical projection neurons typically project to a variety of subcortical targets in the brain, while upper layer neurons (layer II-IV) project to targets within the cortex. The distinctive connection patterns and functional properties of neurons that occupy each cortical layer is essential for the complex circuit computations and behaviors mediated by the cerebral cortex. The extent to which glial cell types which include astrocytes, oligodendrocytes and microglia also have layer-specific molecular and functional properties to support cortical layer specific information processing has not been completely determined (Hammond et al. 2019, Marisca et al. 2020, Pestana et al. 2020).

Astrocytes are the most abundant glial cell type in the CNS and are responsible for supporting a diverse range of functions necessary for cortical circuit activity including neurotransmitter reuptake, ion homeostasis, synapse formation and elimination (Ben Haim and Rowitch 2017). It is unclear whether these diverse functions are mediated by a single class of protoplasmic astrocytes or if there are distinct cell subtypes with specialized roles. Recent studies have shown that protoplasmic astrocytes display differences in their morphological and molecular properties that closely correspond to the organization of cortical projection neurons, suggesting the existence of astrocytes with specialized layer specific functional roles (Batiuk et al. 2020, Bayraktar et al. 2020, Lanjakornsiripan et al. 2018). For example, chordin-like 1 (Chrdl1) an astrocyte secreted factor involved in promoting glutamatergic synapse maturation by inducing the insertion of GluA2 receptors, is differentially enriched in upper layer astrocytes (Blanco-Suarez et al. 2018), while the secreted factor Norrin, which regulates dendritic spine development and electrophysiological properties of neurons is enriched in layer V astrocytes (Miller et al. 2019).

How diverse astrocyte subtypes are specified during development is not understood (Akdemir, Huang and Deneen 2020). Fate mapping suggests that cortical astrocytes derived from the same pool of progenitors display tremendous heterogeneity in their location and morphological features (Clavreul et al. 2019). Comprehensive analysis of the molecular diversity of cortical astrocytes has shown that cells with similar molecular features are localized to specific layers (Bayraktar et al. 2020, Lanjakornsiripan et al. 2018). Evidence suggests that neuron-derived cues provide positional information to astrocytes that can regulate specific aspects of their functions (Bayraktar et al. 2020, Lanjakornsiripan et al. 2018). Alterations in the organization or diversity of cortical neurons directly influences the pattern of layer-specific astrocyte gene expression and morphology. The full complement of neuron-derived factors that may regulate the layer-specific molecular identity of astrocytes are currently unknown. In our study we provide evidence that neuron-derived Sonic hedgehog (Shh) is one such factor that is critical to establish and maintain layer-specific cortical astrocyte identities.

Shh is most widely known for its role as a morphogen and mitogen during embryonic patterning of the developing nervous system (Garcia et al. 2018, Lee, Zhao and Ingham 2016). In the classic model of Shh signaling the 12-pass transmembrane domain receptor Patched 1 (Ptch1) represses the activity of the 7-transmembrane domain receptor Smoothened, which is required for activation of Hedgehog signaling. PTCH1 binding to SHH ligand releases the inhibition of SMO leading to activation of downstream targets such as Gli family transcription factors (Lee et al. 2016). Shh is expressed by subcortical projection neurons in cortical layer V, where it has been shown to regulate the specific patterns of intracortical circuitry (Harwell et al. 2012). Shh signaling is active in a subpopulation of mature astrocytes that are localized primarily to the deep layers of the cortex (Garcia et al. 2010, Hill et al. 2019). In the cerebellum and cortex, Shh signaling in astrocytes promotes the expression of the inward rectifying potassium channel Kir4.1, which is critical for regulating potassium buffering and neuronal excitability (Farmer et al. 2016, Hill et al. 2019). In this study we identify the complement of molecular programs driven by Shh signaling in cortical astrocytes and show that neuron-derived Shh is both necessary and sufficient to promote astrocyte morphogenesis and the layer-specific enrichment of genes in astrocytes involved in regulating synapse formation and function. Finally, we show that cell-autonomous activation of Shh signaling in cortical astrocytes can function to promote excitatory synapse formation. Taken together our data suggests a model whereby secretion of Shh by layer V cortical projection neurons coordinates the function of surrounding astrocytes to promote the formation and maintenance of layer specific circuitry.

## RESULTS

### Expression of the Sonic Hedgehog Receptor Ptch1 is Specific to Astrocytes during Cortical Development

Sonic hedgehog (Shh) and its receptor Boc are both known to be expressed exclusively by specific subpopulations of subcortical and intracortical projection neurons, respectively (Harwell et al. 2012). Astrocytes in a number of CNS regions are known to be responsive to Shh signaling (Farmer et al. 2016, Garcia et al. 2010, Hill et al. 2019). The 12-pass transmembrane Shh receptor Ptch1 is known to be required for normal pathway activation during tissue patterning and nervous system development (Rohatgi, Milenkovic and Scott 2007). In order to understand the role of Ptch1 in mediating Shh signaling in the developing cortex, we characterized the temporal and cell type specific pattern of Ptch1 expression with heterozygous Ptch1-lacZ mutant mice, where a lacZ-neo fusion gene was inserted in the place of the start codon and portions of exon1 and the entire exon 2 of the *Ptch1* gene (Goodrich et al. 1997). We quantified the number of βGal+ cells in the cortex between postnatal day 5 (P5) and P60 and observed cells distributed throughout layers I to VI with the largest number these cells located in layer IV (**Figure 1A-1C**). The number of βGal+ cells exhibits a bimodal distribution over development, with a peak at P15 that coincides with peak Shh expression by layer V cortical neurons (**Figure S1A and S1B**) and then a second peak at P60 when a population of low immunofluorescent intensity of βGal+ cells appear. The number of Ptch1 expressing cells in layer I steadily increased between P5-P60. In layers IV and V, the number of βGal+ cells were lowest at P28, which coincides with the downregulation of neuronal Shh (**Figure 1C and S1A**). Our data suggests that expression of both Shh and Ptch1 is dynamic over development, with cortical Shh expression reaching its highest level during the second postnatal week, while Ptch1 maintains strong expression throughout adulthood.

**Figure 1:**
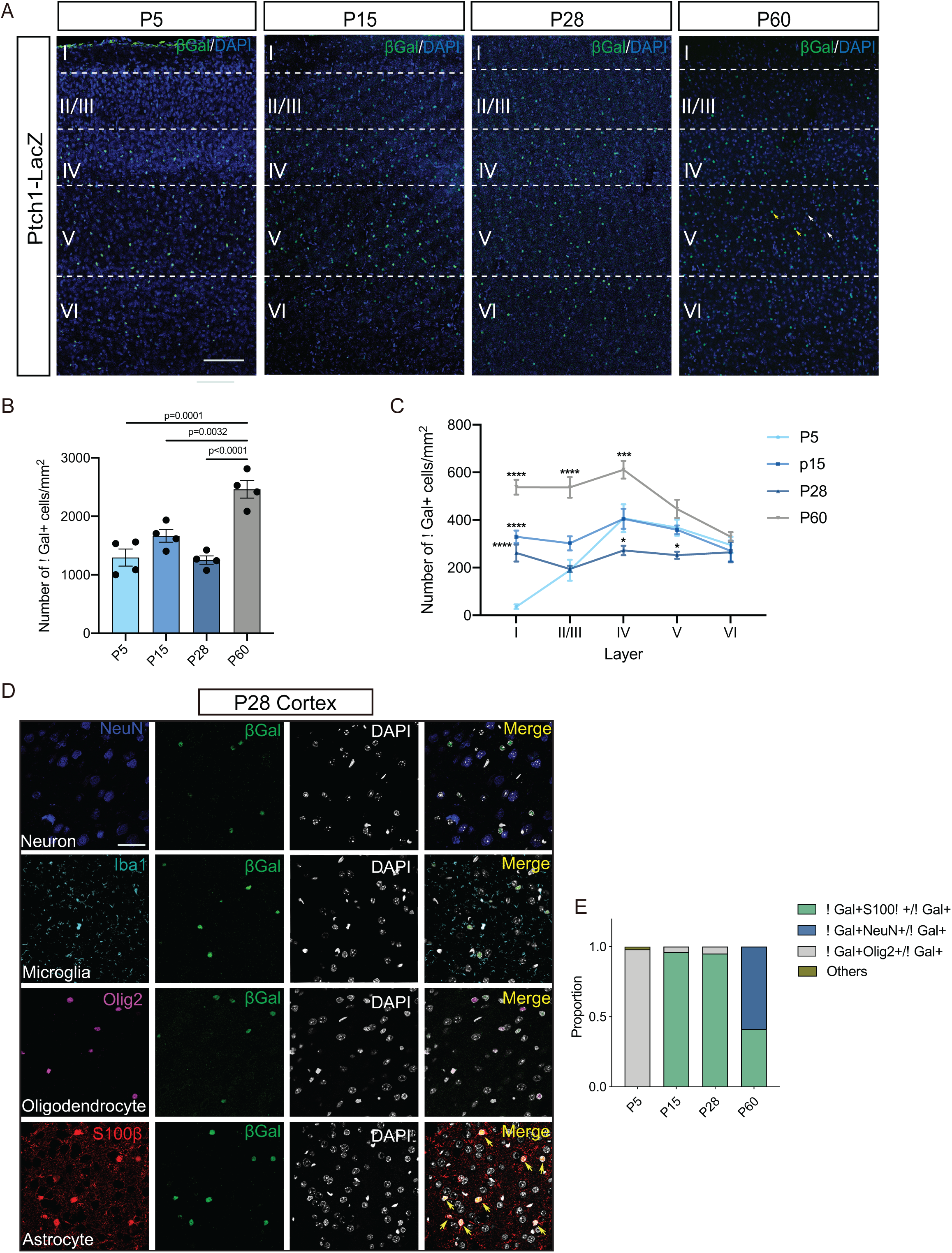
Ptch1 is specifically expressed in cortical astrocytes during development. **A)** Immunostaining for βGal (green) and DAPI (blue) in P5-P60 Ptch1-LacZ mice. Cortical layers are indicated with white dashed lines. White arrows and yellow arrows indicate cells with low and high βGal immunofluorescent intensity, respectively. Scale bar: 250 μm. **B)** Number of βGal+ cells per mm^2^ in entire cortex from P5-P60. N=4 mice for each age. P5 (1296 ± 146); P15 (1666 ± 110.3); P28 (1253 ± 71.05); P60 (2462 ± 150.1). **C)** Number of βGal+ cells per mm^2^ in each cortical layer from P5-P60 Ptch1-LacZ mice. The number of βGal+ cells in P15, P28 and P60 cortex is compared to the number of cells in P5 cortex respectively. N=4 mice for each age. Layer I: P5 (35.94 ± 9.67), P15 (329.72 ± 25.92), P28 (261.48 ± 35.71), P60 (537.82± 31.25); Layer II/III: P5 (189.06 ± 44.06), P15 (302.08 ± 29.90), P28 (194.45 ± 13.20), P60 (536.98 ± 43.50); Layer IV: P5 (407.26 ± 58.28), P15 (404.90 ± 42.03), P28 (272.14 ± 19.82), P60 (611.01 ± 37.03); Layer V: P5 (368.21 ± 33.78), P15 (359.20 ± 17.98), P28 (252.01 ± 15.17), P60 (446.49 ± 38.45); layer VI: P5 (295.06 ± 33.73), P15 (270.09 ± 43.72), P28 (264.60 ± 42.15), P60 (329.50 ± 19.45). **D)** Immunostaining for NeuN (blue), Iba1(cyan), Olig2 (magenta), S100β (red) with βGal (green) and DAPI (grey) respectively in P28 Ptch1-LacZ cortex. Yellow arrows indicate representative S100β+βGal+ cells. Scale bar: 50 μm. **E**) Proportion of cells expressing Ptch1 in P5-P60 cortex. N=3 mice for each age. P5: βGal+Olig2+/βGal+ (98.20%), other cells (1.80%); P15: βGal+S100β+/βGal+ (96%), βGal+NeuN+/βGal+ (0.1%), βGal+Olig2+/βGal+ (3.9%); P28: βGal+S100β+/βGal+ (95%), βGal+NeuN+/βGal+ (0.1%), βGal+Olig2+/βGal+ (4.9%); P60: βGal+S100β+/βGal+ (41%), βGal+NeuN+/βGal+ (59%). Data represent mean ± SEM; statistical analysis is one-way ANOVA with Tukey’s multiple comparisons test (**B**), two-way ANOVA with Dunnett’s multiple comparisons test (**C**). Only p values <0.05 are shown in figures. *p < 0.05, **p < 0.01, and ***p < 0.001; n.s. not significant. See also **Figure S1**.

We further characterized the cell type identities of Ptch1 expressing cells by performing immunofluorescence staining for cell type markers on the cortex of P28 Ptch1-lacZ heterozygous mutant mice. At P5, the majority of βGal+ cells were positive for Olig2, which labels astrocyte and oligodendrocyte precursors at this particular stage. By P15 nearly all βGal+ cells (96%) were positive for the astrocyte marker S100β while few cells (3.9%) were co-labeled with Olig2. At P60, weakly labeled βGal+ cells that were positive for the pan-neuron marker NeuN make up over half of the total βGal+ population (59%) (**Figure 1D and 1E, S1C and S1D**). Taken together, these findings show that Ptch1 is primarily expressed by cortical astrocytes across all layers of the cortex during development, and by P60 it begins to be weakly expressed by subsets of cortical neurons in the mature cortex, suggesting that Shh signaling in the cortex is both cell-type and developmental stage-specific.

### Shh Signaling Influences Astrocyte-specific Transcriptional Programs in Development

Gli1, a direct transcriptional target of Shh pathway activation, is enriched in subgroups of astrocytes located in close proximity to deep layer Shh releasing neurons (Garcia et al. 2010, Allahyari et al. 2019). However, the full complement of genes regulated by activation of Shh signaling in cortical astrocytes is unknown. To address this, we used an astrocyte-specific conditional mutant mouse line, where Shh signaling is hyper-activated by the conditional deletion of Ptch1 (Ptch1cKO). To elevate the level of Shh signaling in astrocytes, we crossed GlastCreER mice with two other lines, one carrying a Cre-recombinase conditional allele of Ptch1 and the other an Ai14 reporter line; GlastCreER drives Cre expression specifically in astrocytes, and labels cells with the fluorescent reporter protein tdTomato after tamoxifen administration at P12-P14 (**Figure 2A**). We sequenced the transcriptomes of FACS purified cortical astrocytes to assess differential gene expression in response to increased levels of Shh signaling (**Figure 2A, S2A and S2B**). We verified the purity of sorted cells by immunofluorescence staining and RT-PCR. TdTomato positive sorted cells showed enriched expression for tdTomato and the astrocyte markers, GLT1 (*Slc1a2*) and *Aldh1l1* (**Figure 2B and S2C**).

**Figure 2:**
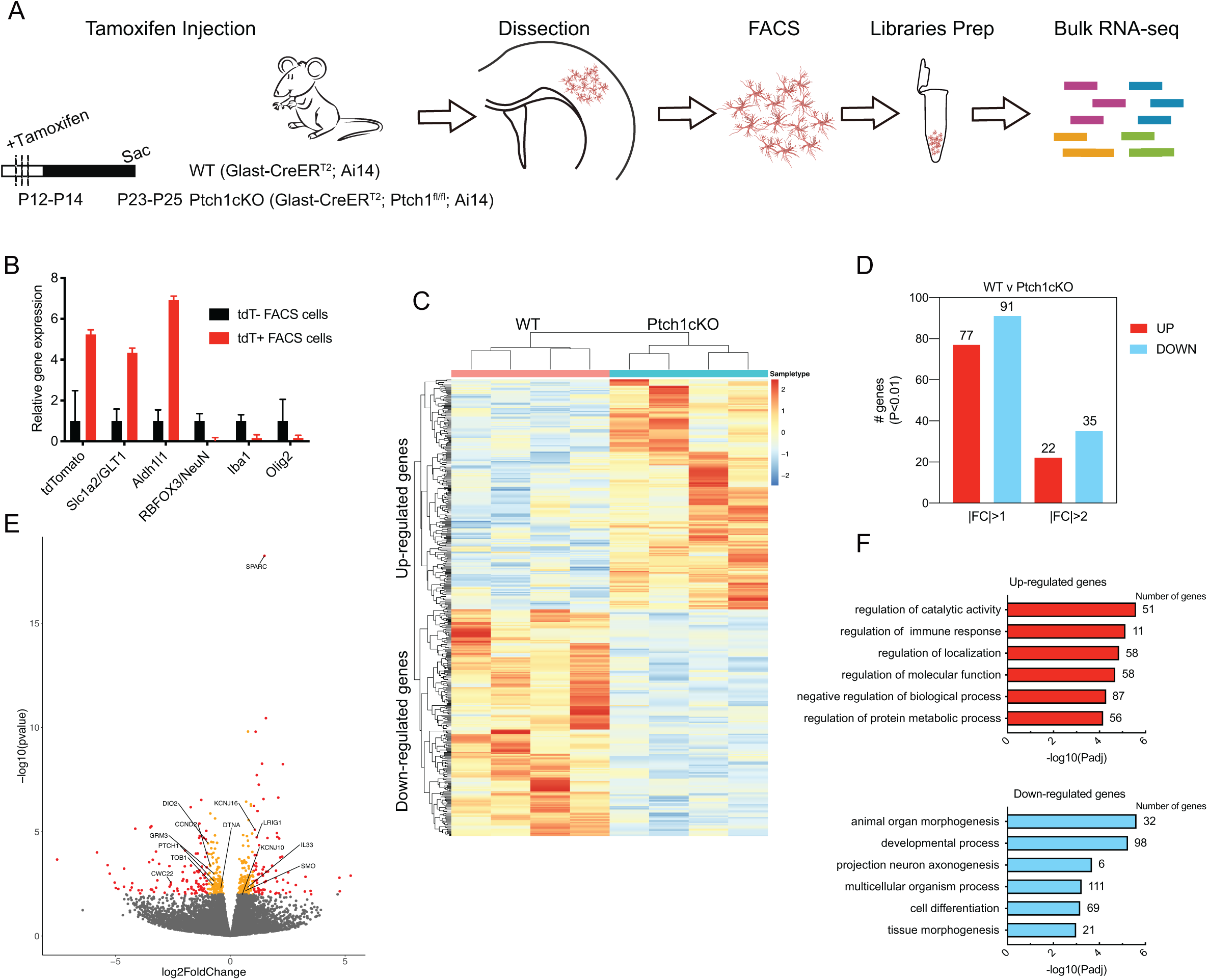
Transcriptional profiling of WT and Ptch1cKO cortical astrocytes during developmental period. **A)** Experimental design for transcriptional profiling of WT and Ptch1cKO cortical astrocytes. **B)** Expression of cell type markers in tdTomato– and tdTomato+ sorted astrocytes. **C**) Heatmap representing changes of gene expression in Ptch1cKO cortical astrocytes compared to WT (p<0.05). **D)** Number of upregulated and downregulated genes in Ptch1cKO vs WT cortical astrocytes (p<0.01). **E)** Volcano plot showing fold changes of gene expression in Ptch1cKO astrocytes. Each dot indicates an upregulated or downregulated gene; red dots (p<0.01, |log2FoldChange|>1); orange dots (p<0.01), remaining dots are grey. **F**) Gene ontology analysis of differentially expressed genes between WT and Ptch1cKO cortical astrocytes. All transcriptome analyses used 4 biological replicates per genotype. See also **Figure S2**.

Differential gene expression analysis revealed consistent changes between WT and Ptch1cKO astrocytes across all four biological replicates (**Figure 2C**). We observed 77 upregulated and 91 downregulated genes in Ptch1cKO astrocytes using |log2FC| > 1 and P value < 0.01 cutoffs (**Figure 2D**). Shh-related genes *Smo* and *Hhip* were significantly increased whereas we observed a 40% reduction of *Ptch1* transcripts in Ptch1cKO compared to WT, suggesting efficient tamoxifen-induced recombination (**Figure S2D**). Functional classification of differentially expressed genes revealed that upregulated genes in Ptch1cKO astrocytes were involved in processes related to cell localization, immune system and cell metabolism, while downregulated genes in Ptch1cKO astrocytes were enriched for GO terms related to nervous system development (**Figure 2F**). Genes that were significantly increased in Ptch1cKO included *Kir4.1*/*Kcnj10, Kir5.1/Kcnj16* (inward rectifying potassium channel), *Lrig1* (leucine-rich repeats and immunoglobulin (Ig)-like domains 1), *Il33* (cytokine interleukin-33) and *Sparc* (secreted protein acidic and rich in cyst) (**Figure 2E**). These genes have previously been shown to be enriched in astrocytes and important factors regulating synapse formation and function (Alsina et al. 2016, Vainchtein et al. 2018, Kucukdereli et al. 2011, Jones et al. 2018). Together this data shows that Shh signaling in cortical astrocytes promotes the expression of genes involved in astrocyte mediated synapse development and circuit function.

### Cell-Autonomous Activation of Shh Signaling in Astrocytes is Sufficient to Promote Layer-specific Gene Expression

Although the bulk sequencing of Ptch1cKO sorted astrocytes provides a picture of the transcriptional programs regulated by Shh signaling in cortical astrocytes, we could not determine whether the transcriptional response to Shh signaling was equivalent across all recombined astrocyte populations in distinct cortical layers. It was also possible that other cell populations apart from astrocytes respond to Shh signaling. We chose to focus our validation on upregulated genes that were specifically enriched in astrocytes. We used RT-PCR and fluorescent *in situ* hybridization to verify the increased expression of *Kir4.1, Lrig1, Sparc* and *Il33* in the cortex of Ptch1cKO mice, and found that all of these genes were significantly increased in cortical astrocytes (**Figure 3D, 3H, 3L, 3P, S4C**), confirming our RNA-seq analysis (**Figure 3C, 3G, 3K, 3O**).

**Figure 3:**
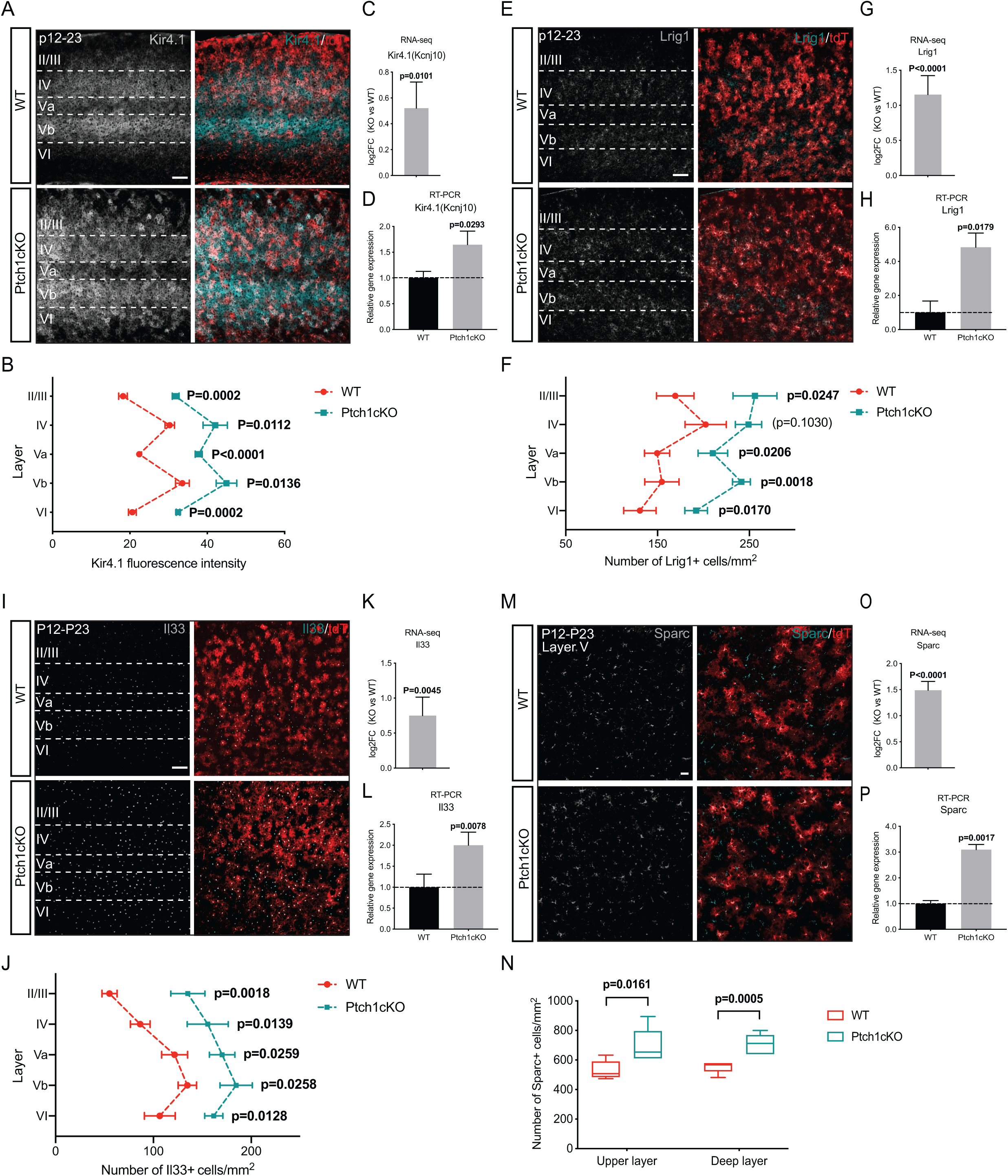
Expression of astrocyte target genes are upregulated in Ptch1cKO. **A)** Mice were injected intraperitoneally with tamoxifen at P12-P14. Immunostaining for Kir4.1(grey/cyan) and tdTomato (red) in P23 WT and Ptch1cKO cortex. Scale bar: 100 μm. **B)** Kir4.1 fluorescence intensity in WT and Ptch1cKO cortex in indicated layers. WT: N=4 mice, KO: N=3 mice. Layer II/III: WT (18.21 ± 1.11), KO (31.83 ± 0.90); layer IV: WT (30.28 ± 1.22), KO (42.06 ± 3.16); layer Va: WT (22.42 ± 0.23), KO (37.80 ± 0.90); layer Vb: WT (33.55 ± 1.78), KO (44.96 ± 2.65); layer VI: WT (20.63 ± 1.03), KO (32.44 ± 0.46). **C, G, K, O)** RNA-seq shows fold-changes of *Kir4.1/Kcnj10* (**C***), Lrig1*(**G**)*, Il33* (**K**) *and Sparc* (**O**) genes expression from Ptch1cKO compared to WT. N=4 mice for each condition. **D, H, L, P**) Realtime PCR shows relative gene expression of *Kir4.1/Kcnj10* (**D**), *Lrig1* (**H**), *Il33* (**L**) *and Sparc* (**P**) from Ptch1cKO compared to WT. N=4 mice for each condition. *Kir4.1*: WT (1.00 ± 0.13), KO (1.57 ± 0.26); *Lrig1*: WT (1.00 ± 1.00), KO (4.83 ± 0.83); *Il33*: WT (1.00 ± 0.31), KO (2.00 ± 0.31); *Sparc*: WT (1.00 ± 0.12), KO (3.10 ± 0.19). **E**) Immunostaining for Lrig1(grey/cyan) and tdTomato (red) in P23 WT and Ptch1cKO cortex. Scale bar: 100 μm. **F**) Number of Lrig1+ cells in WT and Ptch1cKO in indicated layers. WT: N=5 mice, KO: N=6 mice. Layer II/III: WT (169.21 ± 20.41), KO (256.06 ± 23.96); layer IV: WT (202.43 ± 22.51), KO (249.14 ± 14.31); layer Va: WT (149.49 ± 13.67), KO (210.22 ± 16.12); layer Vb: WT (154.68 ± 18.67), KO (241.36 ± 9.73); layer VI: WT (130.80 ± 17.63), KO (192.05 ± 12.36). **I**) Immunostaining for Il33 (grey/cyan) and tdTomato (red) in P23 WT and Ptch1cKO cortex. Scale bar: 100 μm. **J**) Number of Il33+ cells in indicated cortical layers from WT and Ptch1cKO. N=6 mice for each condition. Layer II/III: WT (55.129 ± 7.742), KO (135.16 ± 17.44); layer IV: WT (86.57 ± 10.03), KO (155.58 ± 20.93); layer Va: WT (121.69 ± 13.28), KO (170.28 ± 13.03); layer Vb: WT (134.76 ± 9.42), KO (184.58 ± 16.56); layer VI: WT (106.58 ± 15.73), KO (161.71 ± 9.23). **M**) Immunostaining for Sparc (grey/cyan) and tdTomato (red) in P23 WT and Ptch1cKO layer V cortex. Scale bar: 30 μm. **N**) Number of Sparc+ cells in upper and deep layers from WT and Ptch1cKO cortex. N=6 mice for each condition. Deep layer: WT (551.599 ± 15.46), KO (710.15 ± 27.74); upper layer: WT (530.58 ± 25.16), KO (693.43 ± 52.76). Data represent mean ± SEM; statistical analysis is multiple t-test. See also **Figure S3 and S4.**

Next we performed immunohistochemistry to determine the cell type- and layer-specific expression patterns of Shh target genes. Previous work has shown that Kir4.1 is expressed exclusively in glial cells in the CNS, and is thought to be required for potassium buffering in the brain (Djukic et al. 2007, Tong et al. 2014). We found that Kir4.1 expression was highly enriched in cortical layer IV and layer Vb WT astrocytes, which extensively overlaps with the location of Shh expressing neurons (**Figure 3A, S1A**). In Ptch1cKO mice, Kir4.1 fluorescence intensity was dramatically increased across all layers compared to WT (**Figure 3A and 3B**), suggesting that astrocytes across all layers of the cortex have a similar capacity to respond to Shh signaling. We also observed increased Shh-dependent expression of Kir5.1, a pH sensitive inward rectifying potassium channel that is capable of forming heteromeric complexes with Kir4.1 (Brasko et al. 2017) (**Figure S4A and S4B**). Next we verified whether the expression of Lrig1, Il33 and Sparc was increased in response to elevated levels of Shh signaling. In WT animals, we observed that Lrig1 was most highly expressed in layer IV, largely overlapping with tdTomato+ astrocytes at P21 (**Figure 3E**), while Il33 was predominantly expressed in deep layer astrocytes (**Figure 3I**), consistent with previous analyses suggesting this gene may serve as a deep layer astrocyte marker (Bayraktar et al. 2020). Sparc+ cells were evenly distributed in between upper and deep layers (**Figure S3A**) and found in both tdTomato+ astrocytes (**Figure 3M**) and Iba1+ microglia cells (**Figure S3B**). As expected, we observed significantly increased numbers of Lrig1+, Il33+ and Sparc+ cells throughout upper and deep layers of Ptch1cKO cortex (**Figure 3E, 3F, 3I, 3J, 3M and 3N**). However, we did not observe an increase in the number of Iba1+Sparc+ cells in Ptch1cKO (**Figure S3B and S3C**). Our findings indicate that Shh signaling is sufficient to promote the expression of layer-specific genes in cortical astrocytes during development.

### Sonic Signaling is Necessary for the Expression of Layer-specific Astrocyte Genes

In order to test whether layer V neuron-derived Shh was specifically required to maintain the expression of target genes in cortical astrocytes, we generated a mouse line where Shh is specifically deleted in the cortex (ShhcKO) by crossing the Emx1 *^ires^*^-cre^ mouse line with a line carrying a conditional *Shh* deletion allele (Shh^flox/flox^). We analyzed the abundance and distribution of Kir4.1 in the cortex using immunohistochemistry in P21 WT and ShhcKO mice. We found that Kir4.1 expression was significantly decreased in layers IV and Vb of ShhcKO mice at P21 (**Figure 4A-4C**). This is consistent with previously reported observations where disrupted Shh signaling by deletion of the Shh receptor Smo led to reduced Kir4.1 expression (Hill et al. 2019, Farmer et al. 2016). In order to investigate whether Kir5.1 was also decreased in ShhcKO cortical neurons, we used single molecule fluorescent *in situ* hybridization to detect *Kir5.1* transcripts and observed a similar pattern of significantly reduced *Kir5.1* expression in the deep layers of cortex (**Figure S4D, S4E**).

**Figure 4:**
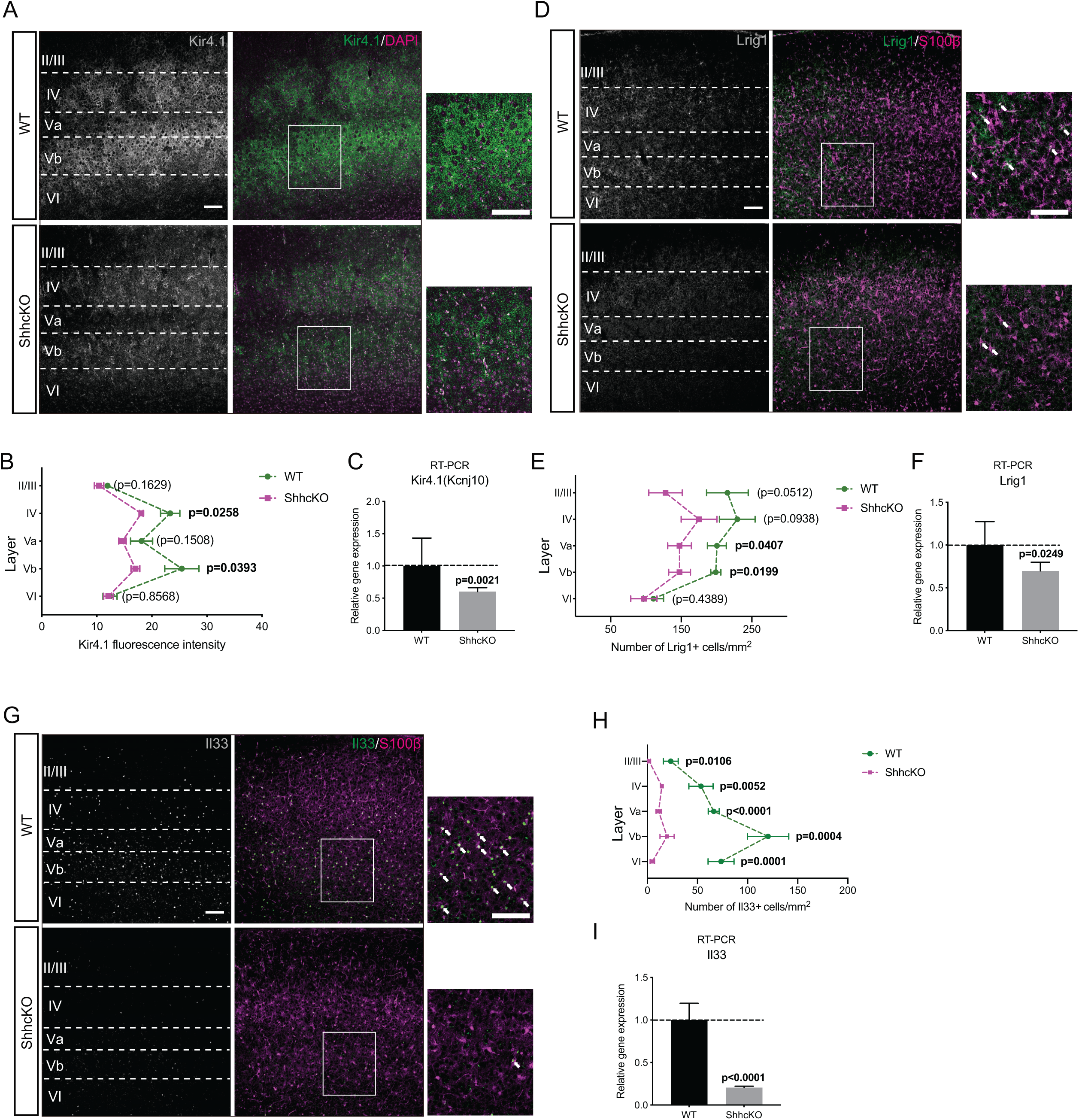
Expression of target genes is downregulated in ShhcKO. **A**) Immunostaining for Kir4.1(grey/green) and DAPI (magenta) in P21 WT and ShhcKO cortex. Cortical layers are indicated in white. High magnification images show Kir4.1 is highly enriched in layer V cortex. Scale bar: 100 μm. **B**) Kir4.1 fluorescence intensity in WT and ShhcKO in indicated layers. N=4 mice for each condition. Layer II/III: WT (11.94 ± 0.43), KO (10.41 ± 0.86); layer IV: WT (23.33 ± 1.75), KO (18.07 ± 0.35); layer Va: WT (18.13 ± 2.00), KO (14.67 ± 0.67); layer Vb: WT (25.44 ± 3.11), KO (17.01 ± 0.81); layer VI: WT (12.40 ± 1.28), KO (12.11 ± 0.88). **C**, **F**, **I**) Realtime PCR shows relative gene expression of *Kir4.1/Kcnj10* (**C**), *Lrig1* (**F**), *Il33* (**I**) from ShhcKO vs WT. *Kir4.1*: WT (N=5 mice, 1.00 ± 0.43), KO (N=5 mice, 0.60 ± 0.06); *Lrig1*: WT (N=5 mice, 1.00 ± 0.27), KO (N=5 mice, 0.70 ± 0.10); *Il33*: WT (N=5 mice, 1.00 ± 0.20), KO (N=5 mice, 0.20 ± 0.02); **D**) Immunostaining for Lrig1(grey/green) and S100β (magenta) in P21 WT and ShhcKO cortex. Cortical layers are indicated in white. High magnification images show Lrig1 is enriched in deep layer S100β+ astrocytes. White arrows indicate representative Lrig1+S100β+ cells. Scale bar: 100 μm. **E**) Number of Lrig1+ cells in WT and ShhcKO in indicated layers. WT: N=6 mice, KO: N=5 mice. Layer II/III: WT (215.36 ± 29.48), KO (127.73 ± 23.71); layer IV: WT (229.39 ± 25.17), KO (175.26 ± 25.37); layer Va: WT (200.51 ± 13.45), KO (147.54 ± 16.82); layer Vb: WT (198.86 ± 7.17), KO (147.54 ± 15.45); layer VI: WT (110.57 ± 14.27), KO (97.04 ± 18.56). **G**) Immunostaining for Il33 (grey/green) and S100β (magenta) in P21 WT and ShhcKO cortex. Cortical layers are indicated in white. High magnification images show Il33 expression in deep layer astrocytes. White arrows indicate representative Il33+S100β+ cells. Scale bar: 100 μm. **H**) Number of Il33+ cells in indicated cortical layers from WT and ShhcKO. WT: N=6 mice, KO: N=7 mice. Layer II/III: WT (23.28 ± 7.45), KO (1.75 ± 1.38); layer IV: WT (53.49 ± 12.07), KO (14.35 ± 1.88); layer Va: WT (66.15 ± 5.44), KO (11.20 ± 2.55); layer Vb: WT (120.46 ± 20.62), KO (19.60 ± 7.09); layer VI: WT (73.50 ± 12.98), KO (4.90 ± 2.13). Data represent mean ± SEM; statistical analysis is t-test or multiple t-test. See also **Figure S4 and S5**.

We also detected significant reductions of both protein and RNA for the layer-specific astrocyte enriched genes *Lrig1* and *Il33* (**Figure 4D-4I, S4F**), as well as a non-layer-specific gene *Sparc,* in the cortex of ShhcKO mice (**Figure S4G-S4I**). Surprisingly, we observed that the number of Il33+ cells was comparable to control levels in both upper and deep layers of adult ShhcKO mice (**Figure S5A and S5B**). Previous research has shown that in the adult mouse CNS, Il33 is mainly expressed by oligodendrocytes (Gadani et al. 2015). We immunostained the cortex of 4-month-old WT mice for Il33 and the oligodendrocyte marker Olig2 and found that Il33 was primarily enriched in Olig2+ oligodendrocytes (**Figure S5C**). Thus, the expression pattern of Il33 appeared to shift from Shh-signaling-dependent astrocyte-specific to Shh-independent oligodendrocyte-specific expression over the course of development. Collectively, this data shows that secretion of Shh by layer V cortical neurons is required to maintain expression of layer-specific astrocyte genes during development.

### Ectopic Neuronal Expression of Shh Promotes Target Genes Expression in Local Astrocytes

Recent studies have shown that the morphological and molecular differences between cortical astrocytes are driven by neuronal layers (Bayraktar et al. 2020, Lanjakornsiripan et al. 2018). To determine if a neuron-specific source of Shh is sufficient to induce astrocytes to adopt a deep layer astrocyte molecular profile, we mis-expressed a full length Shh transgene in upper layer neurons, where it is typically absent. We co-electroporated plasmids encoding pCAG-IRES-Cre and Flex-Shh-IRES-GFP into the lateral ventricles of embryonic day (E)15 mouse embryos, a period when the majority of cortical progenitors generate upper layer neurons. Plasmids encoding pCAG-IRES-Cre and Flex-tdTomato were used as controls. Brains were collected at P14 and stained for Kir4.1, Lrig1, Il33 and Sparc. We did not observe any differences in the survival or migration of upper layer neurons overexpressing Shh compared to electroporations with control plasmids. Compared to the non-electroporated contralateral side and control plasmid electroporations, astrocyte-specific target genes expression was increased in upper layer cortical astrocytes in proximity to the somatodendritic region of neurons overexpressing Shh (**Figure 5A-5H**). These results suggest that Shh can be released from the somatodentritic region of neurons and is sufficient to promote deep layer astrocyte transcriptional programs in the developing cortex.

**Figure 5:**
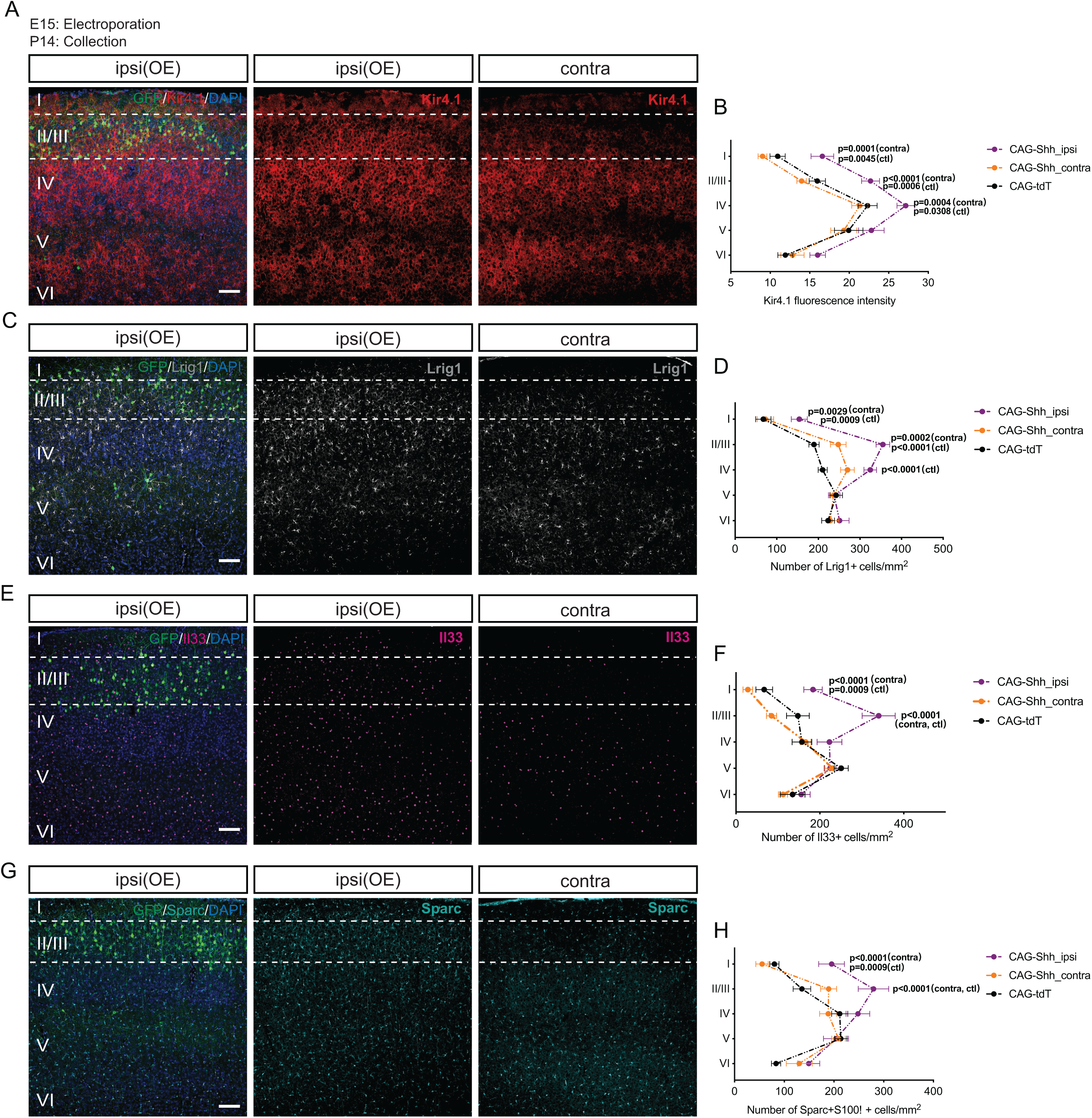
Ectopic expression of Shh in upper layer neurons increases target genes expression in astrocytes. **A-H**) Immunostaining for Kir4.1 (red) (**A**), Lrig1 (grey) (**C**), Il33 (magenta) (**E**), Sparc (cyan) (**G**) combined with GFP (electroporated cells) and DAPI (blue) respectively in P14 mice electroporated at E15 with control or Shh overexpression plasmids (See **Key Resources Table**). Cortical layers are indicated in white. Scale bar: 100 μm. There is an increase in Kir4.1 fluorescence intensity (**B**), number of Lrig1+ cells (**D**), number of Il33+ cells (**F**) and number of Sparc+S100β+ cells (**H**) in indicated layers from electroporated hemispheres (ipsilateral, ipsi; purple) compared to non-electroporated contralateral hemispheres (contra; orange) and electroporated controls (ctl; black). N=5-7 mice for each group, Kir4.1: layer I (ipsi: 16.59 ± 1.43, contra: 9.016 ± 0.56, ctl: 10.94 ± 0.97), layer II/III (ipsi: 22.70 ± 1.13, contra: 13.97 ± 0.60, ctl: 15.96 ± 1.03), layer IV (ipsi: 27.16 ± 1.11, contra: 21.34 ± 1.02, ctl: 22.32 ± 1.20), layer V (ipsi: 22.80 ± 1.62, contra: 19.32 ± 1.65, ctl: 19.93 ± 1.81), layer VI (ipsi: 16.00 ± 0.99, contra: 12.82 ± 1.47, ctl: 11.91 ± 0.96). Lrig1: layer I (ipsi: 154.24 ± 19.23, contra: 72.87 ± 19.69, ctl: 68.01± 18.31), layer II/III (ipsi: 355.15 ± 16.36, contra: 260.40 ± 19.02, ctl: 189.51 ± 12.21), layer IV (ipsi: 324.77 ± 15.21, contra: 285.13 ± 19.59, ctl: 210.49 ± 11.02), layer V (ipsi: 237.97 ± 13.70, contra: 236.53 ± 7.55, ctl: 243.04 ± 15.20), layer VI (ipsi: 250.93 ± 22.73, contra: 228.71 ± 20.92, ctl: 223.50 ± 15.23). Il33: layer I (ipsi: 183.88 ± 22.13, contra: 28.36 ± 11.42, ctl: 67.49 ± 19.90), layer II/III (ipsi: 340.41 ± 39.57, contra: 84.92 ± 11.88, ctl: 147.68 ± 27.07), layer IV (ipsi: 223.00 ± 29.64, contra: 166.15 ± 12.78, ctl: 157.28 ± 23.33), layer V (ipsi: 224.48 ± 21.36, contra: 229.65 ± 17.43, ctl: 251.06 ± 16.50), layer VI (ipsi: 155.81 ± 21.36, contra: 112.24 ± 10.80, ctl: 135.13 ± 29.47). Sparc: layer I (ipsi: 195.29 ± 25.91, contra: 55.38 ± 12.54, ctl: 80.16 ± 9.41), layer II/III (ipsi: 279.49 ± 30.65, contra: 189.22 ± 16.16, ctl: 135.62 ± 17.75), layer IV (ipsi: 248.24 ± 23.91, contra: 188.35 ± 17.28, ctl: 211.57 ± 16.28), layer V (ipsi: 203.98 ± 22.03, contra: 130.20 ± 26.00, ctl: 83.54 ± 9.27), layer VI (ipsi: 149.29 ± 22.03, contra: 130.20 ± 26.00, ctl: 83.54 ± 9.27). Data represent mean ± SEM; statistical analysis is two-way ANOVA with Turkey’s multiple comparisons test. Only p values <0.05 are shown in figures.

### Shh Signaling is required for Astrocyte Morphogenesis and Coverage of Neuronal Synapses

Protoplasmic astrocytes have complex arbors containing fine cellular processes that infiltrate in the neuropil and interact with synapses (Bushong et al. 2002, Allen and Eroglu 2017). Astrocytes that occupy different cortical layers exhibit diverse morphological features (Lanjakornsiripan et al. 2018). We investigated whether neuron-derived Shh expression is required for astrocytes to acquire their unique morphological properties. We utilized postnatal astrocyte labeling by electroporation (PALE) (Stogsdill et al. 2017) with a plasmid encoding membrane *tdTomato* and nuclear GFP (pCAG-mTdT-2A-H2BGFP) to sparsely label astrocytes across the entire cortex of WT and ShhcKO mice (**Figure 6A and 6B**). Sholl analysis of fluorescent labeled astrocytes showed that deep layer astrocytes have a significant reduction in morphological complexity in ShhcKO mice compared to controls, whereas we observed no change in upper layer astrocytes (**Figure 6C and 6D**). Loss of cortical Shh did not lead to a reduction in the number or altered distribution of S100β+ astrocytes (**Figure S6A and S6B**). We next tested whether activation of Shh in cortical astrocytes might yield increased morphological complexity in tdTomato+ Ptch1cKO astrocytes when compared to control labeled cells. Sholl analysis did not reveal significant differences in morphological complexity between Ptch1cKO and controls (**Figure S6C and S6D**). Taken together, our data suggests that Shh signaling is necessary to maintain morphological properties of layer-specific astrocytes *in vivo*.

**Figure 6:**
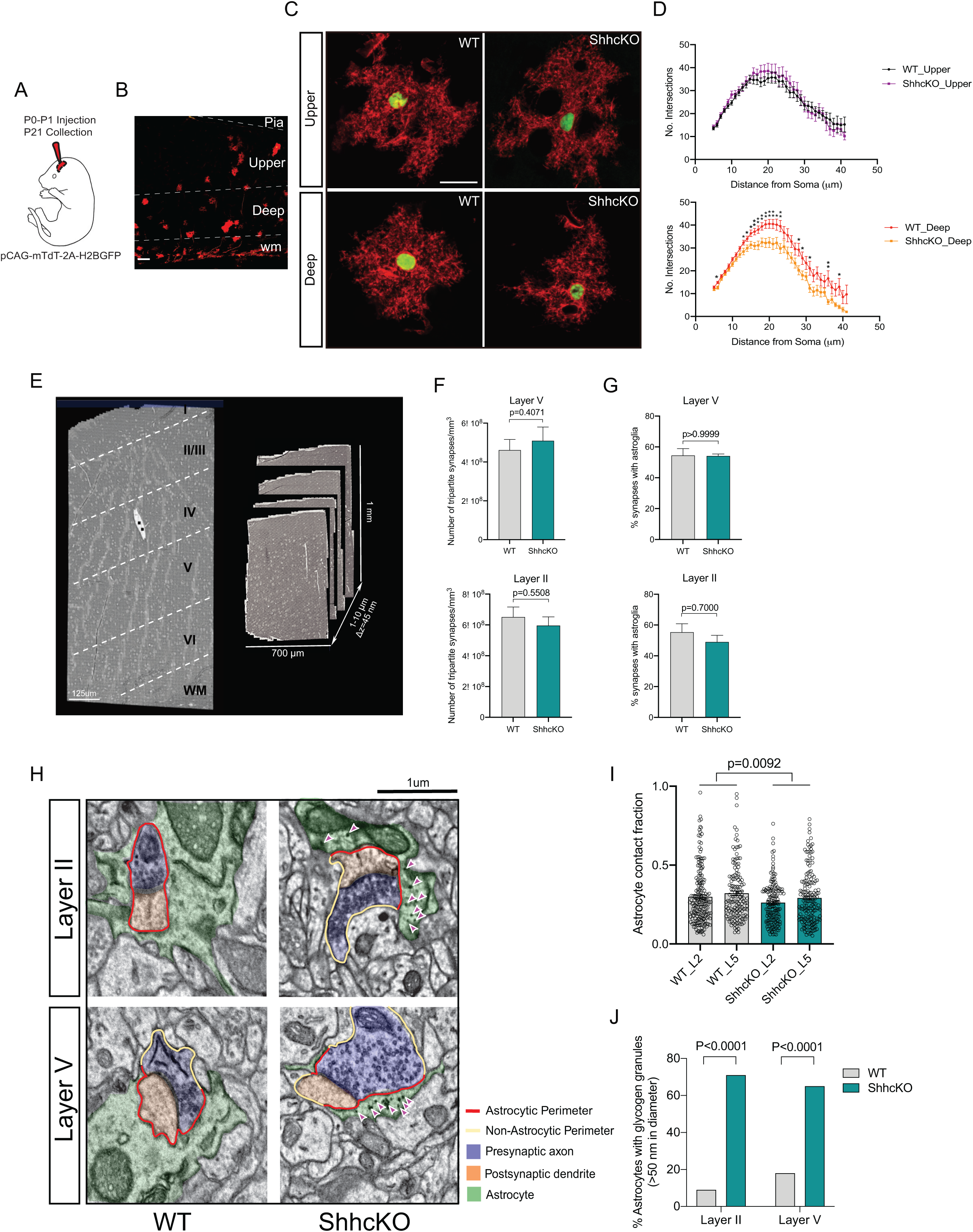
Deletion of Shh reduces astrocyte morphological complexity and coverage of neuronal synapses. **A**) Schematic of Postnatal Astrocyte Labeling by Electroporation with plasmid DNA (pCAG-mTdT-2A-H2BGFP) (See **Key Resource Table**) in early postnatal mice. Samples were collected at P21. **B**) Representative image of P21 cortex following electroporation. TdTomato (processes) and GFP (nuclei) labelled astrocytes are dispersed throughout all layers. Scale bar: 100 μm. **C**) Representative high-magnification confocal images of P21 PALE astrocytes from upper and deep layers of WT and ShhcKO. Scale bar: 20 μm. **D**) Quantification of astrocyte complexity for deep (WT: red; KO: orange) and upper (WT: black; KO: purple) astrocytes. Deep: WT (N=4 mice, n=35 cells), KO (N=4 mice, n=32 cells), upper: WT (N=4 mice, n=35 cells), KO (N=4 mice, n=25 cells). **E**) A large-scale electron microscope image with serial sections from P26 WT mice. Cortical layers are indicated in white. The size of the serial imaging block was 700 μm (X) × 1mm (Y) × (1-10) μm (Z). The thickness of individual images was 45 nm. Scale bar: 125 μm. **F**) Number of tripartite synapses per mm^3^ is not significantly different in either layer II or layer V of ShhcKO compared to WT. Layer V: WT (4.61 × 10^8^ ± 3.16 × 10^7^), KO (5.09 × 10^8^ ± 4.09 × 10^7^). Layer II: WT (6.52 × 10^8^ ± 6.43 × 10^7^), KO (5.96 × 10^8^ ± 5.61 × 10^7^). **G**) Proportion of tripartite synapses is not significantly changed in either layer II or layer V of ShhcKO cortex. Layer V: WT (54.49 ± 2.55), KO (54.17 ± 0.73). Layer II: WT (55.39 ± 5.45), KO (49.02 ± 4.27). **H**) Representative EM images acquired from layer II and layer V cortex of WT and ShhcKO. Magenta arrows indicate dark glycogen granules in astrocytic processes. Scale bar: 1 μm. **I**) Quantification of astrocyte ensheathment of synapses in layer II and layer V cortex of WT and ShhcKO. Layer II: WT (n=194 synapses), KO (n=162 synapses); Layer V: WT (n=142 synapses), KO (n=166 synapses). **J**) Proportion of astrocytes with intensely labeled glycogen granules (>50 nm in diameter). Layer II: WT (n=18/194), KO (n=112/162); Layer V: WT (n=28/142), KO (n=109/166). Data Represent mean ± SEM; statistical analysis is multiple t-test (**D**), Welch’s test (**F**) and Mann-Whitney test (**G**), two-way ANOVA test (**I**) and Chi-squared test (**J**). See also **Figure S6, Table S1 and Table S2**.

Astrocytes are intimately associated with surrounding neuronal synapses to form tripartite synapses (Araque et al. 1999, Papouin et al. 2017, Farhy-Tselnicker and Allen 2018). In order to test the possibility that Shh signaling might influence the formation of perisynaptic astrocyte processes, we utilized electron microscopy (EM) to examine the structure and connectivity of astrocyte associated synapses across layers of the cortex. We generated EM image volumes (Graham et al. 2019) spanning all six layers of the somatosensory cortex of WT and ShhcKO animals (**Figure 6E**) and quantified structural features of synapses. We first quantified the density of synapses contained within three randomly positioned boxes (20×10×0.9 μm) in layer II and layer Vb of P26-P28 WT and ShhcKO samples. Volumes representative of layer II and layer V were chosen at the same respective distance from the corpus callosum in both WT and ShhcKO samples. Synapses were identified by their characteristic structural features, namely synaptic vesicles in the presynaptic compartments along with darkly stained postsynaptic densities. We also counted the number of perisynaptic astrocyte processes in direct contact with the synaptic cleft within an imaging volume known as tripartite synapses. Astrocytes processes were distinguished by their irregular shape and dense glycogen granules (Cali et al. 2019a, Kikuchi et al. 2020). We manually traced 2244 synapses and their associated astrocyte processes (456 WT layer V, 496 KO layer V, 636 WT layer II, 656 KO layer II) across 20 well-aligned serial sections. 3D reconstructed images from WT and ShhcKO show astrocyte processes were surrounding synapses at the level of synaptic cleft (**Figure 6SE**). We did not detect a change in the number of tripartite synapses in either layer II or layer V of ShhcKO cortex compared to controls (**Figure 6F)**, or a change in the number of neuronal synapses (**Figure S6F**). We also did not observe a change in the proportion of synapses that were associated with astrocyte processes (**Figure 6G**). We further investigated whether Shh signaling might influence the structure of perisynaptic astrocyte processes by measuring the extent of ensheathment of synaptic clefts by astrocytes in layer II and layer V. Intriguingly, our data showed that the fraction of synaptic interface perimeter that was surrounded by perisynaptic astroglia in layer V KO was significantly reduced compared to WT (**Figure 6H,6I and Table S1**), suggesting that these astrocytes in ShhcKO have less structural connection with surrounding deep layer neurons in ShhcKO. We also observed more astrocytes with intensely labeled glycogen granules accumulated in their processes compared to WT (**Figure 6H, 6J, S6G and Table S2**). Accumulation of glycogen granules has been shown correlate with astrocytes in reactive states (Cai et al. 2020, Dienel and Rothman 2019, Cali, Tauffenberger and Magistretti 2019b). We also observed significantly increased GFAP labeling in astrocytes in all cortical layers of ShhcKOs (**Figure S6H and S6I**), suggesting that astrocytes become reactive when Shh is deleted in cortical neurons. Together our data suggests that Shh signaling in the cortex is necessary for astrocyte morphogenesis and proper formation of astrocyte-neuron structural connections.

### Activation of Shh Signaling in Astrocytes Promotes Cortical Synapse Formation

Astrocytes are viewed as integral elements of excitatory synaptogenesis and our molecular analysis suggests that Shh signaling promotes the expression of genes involved in synaptic function (Baldwin and Eroglu 2017). To determine whether activation of Shh signaling in astrocytes might function to promote synaptogenesis, we quantified the synapse density within the territory of WT and Ptch1cKO tdTomato astrocytes. Synapses were labelled by the co-localization of pre- and post-synaptic markers: VGluT1/PSD95 (V1) and VGluT2/PSD95 (V2), that represent cortical excitatory and thalamocortical excitatory inputs, respectively. Automated counting of colocalized puncta was performed using MATLAB (**Figure S7A**). We found the density of both V1 and V2 colocalized puncta was significantly increased within Ptch1cKO astrocyte domains in both layer II and layer V (**Figure 7A-7D**). However, deletion of Shh decreased the density of V2 colocalized puncta in deep layer PALE territory, but did not significantly reduce the density of V1 colocalized puncta (**Figure S7B and S7C**). This could be attributable to the diversity of VGluT1+ corticocortical inputs, and their capacity to compensate for Shh-dependent connections.

**Figure 7:**
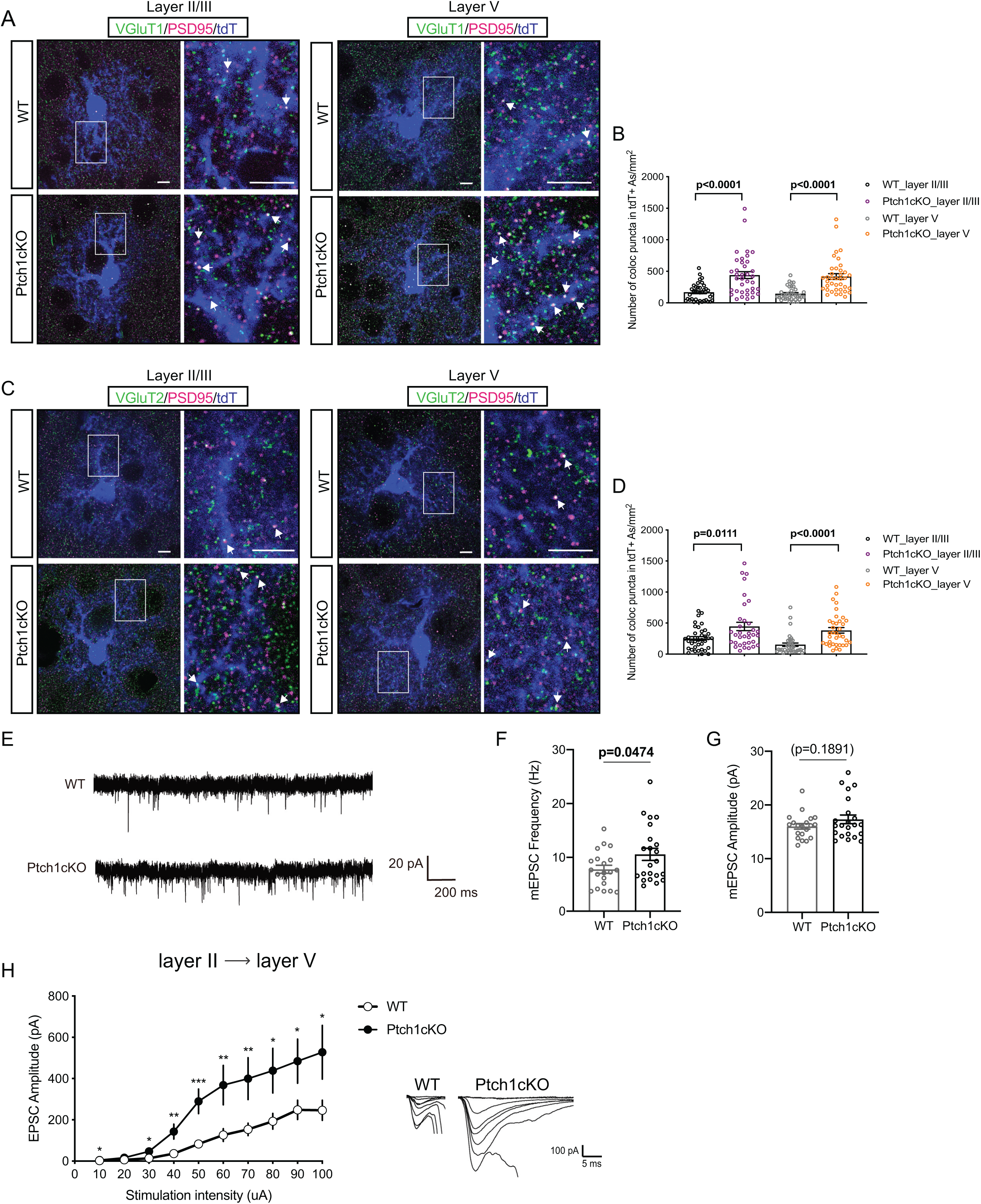
Activation of Sonic signaling in astrocytes promotes cortical synaptogenesis. Mice were injected intraperitoneally with tamoxifen between P12-14 and harvested at P21. (**A**) Immunostaining for intracortical (VGluT1 (green)/ PSD95 (magenta)) (**C**) and thalamocortical (VGluT2 (green)/ PSD95 (magenta)) excitatory synapses in layers II/III and layer V of P21 WT and Ptch1cKO, Astrocytes are labelled with tdTomato (blue). High magnification images of highlighted areas are shown in right panels. White arrows indicate VGluT1/PSD95 and VGluT2/PSD95 colocalized puncta in astrocytes territory. Scale bar: 5 μm. **B**, **D**) Quantification of colocalized puncta density in tdTomato+ astrocytes domains per mm^2^. N=4 mice for each condition. VGluT1/PSD95: Upper layer: WT (n= 37 cells, 168.63 ± 22.51), KO (n=37 cells, 440.45 ± 54.07); deep layer: WT (n=37 cells, 145.65 ± 16.84), KO (n=38 cells, 416.98 ± 45.30). VGluT2/PSD95: Upper layer: WT (n= 37 cells, 252.7 ± 30.24), KO (n=34 cells, 446.5 ± 66.66); deep layer: WT (n=36 cells, 151.9 ± 26.89), KO (n=36 cells, 383.00 ± 44.27). **E**-**G**) Recordings of mEPSCs from layer V neurons shows a significant increase in mEPSCs frequency of Ptch1cKO cells compared to WT, while there is no significant change in mEPSCs amplitude. N=3 mice for each condition (WT: n=20 cells, KO: n=22 cells). mEPSC Frequency: WT (7.79 ± 0.75), KO (10.57 ± 1.13). mEPSC amplitude: WT (16.02 ± 0.52), KO (17.32 ± 0.82). **H**) Evoked EPSC amplitude is significantly increased in layer V neurons of Ptch1cKO cortex when stimuli of different intensities are applied to layer II cortex. N=4 mice for each condition (WT: n=20 cells, KO: 15 cells). Data represent mean ± SEM; statistical analysis is Welch’s t test (**B**, **D**, **F**, **G**) or multiple t test (**H**). *p < 0.05, **p < 0.01, and ***p < 0.001; n.s., not significant. See also **Figure S7**.

We asked whether increased glutamatergic synapses in Ptch1cKO leads to detectable changes in functional circuitry. We performed whole-cell voltage clamp recording and examined spontaneous miniature excitatory postsynaptic currents (mEPSCs) in layer V pyramidal neurons in acute brain slices from P21-P28 WT and Ptch1cKO mice. We found that frequency of spontaneous mEPSCs in layer V neurons was increased (35.75%) in Ptch1cKO slices, while there was no change in mEPSC amplitude, input resistance or membrane capacitance (**Figure 7E-7G, S7D and S7E**). To further determine whether cell-autonomous activation of Shh signaling in cortical astrocytes promotes layer II to layer V connectivity, we examined the evoked EPSCs in layer V pyramidal neurons which was induced by a serial electrical stimulation of layer II neurons. Layer V neurons in Ptch1cKO showed significantly increased EPSC amplitude in response to layer II stimulation (**Figure 7H**). Taken together, our data shows that astrocyte specific activation of Shh signaling promotes the formation of glutamatergic synapses and enhances the strength of layer II to layer V connections.

## DISCUSSION

In this study we identify neuron-derived Shh as a critical factor for organizing the molecular and functional features of deep layer cortical astrocytes. Shh signaling regulates the expression of a diverse repertoire of deep layer astrocyte-enriched target genes with known roles in regulating various aspects of synaptic transmission including, synapse formation (*Sparc, Hevin/Sparcl1, Lrig1*) (Alsina et al. 2016, Jones et al. 2011, Singh et al. 2016), refinement (*Il33*) (Vainchtein et al. 2018), ion homeostasis and neurotransmitter uptake (*Kir4.1*/*Kcnj10 and Kir5.1*/*Kcnj16*) (Brasko et al. 2017, Djukic et al. 2007). The diversity of molecular functions of Shh target genes suggests that signaling shapes multiple aspects of astrocyte development and function, impacting morphology, the secretion of synaptogenic factors and cellular adhesions (Allen and Eroglu 2017). We show that increased Shh signaling in astrocytes promotes the formation of both intracortical and thalamocortical excitatory synapses in the cortex (**Figure 7**). However, the contribution of individual target genes to the increase of specific types of synapses is not clear. Previous research has shown that individual Shh regulated genes such as *Sparcl1* can differentially impact the formation of thalamocortical synapses while having no effect on intracortical connections (Risher et al. 2014). It is also possible that Shh target genes in deep layer astrocytes may indirectly affect synapses, through the regulation of neuron metabolism and ion homeostasis. Shh is expressed by layer Vb subcortical projection neurons that have distinctive intrinsic membrane and firing properties and may require specialized support from local astrocytes (Hattox and Nelson 2007). The inward rectifying potassium channel Kir4.1 has been shown to control the growth of spinal cord motor neurons and burst firing activity of neurons in the lateral habenula that control depression-like behaviors (Cui et al. 2018, Kelley et al. 2018). It is possible that Shh dependent expression of Kir4.1 and Kir5.1 in deep layer astrocytes may support the unique functional properties of layer Vb subcortical projections in a similar way. In addition to genes with known roles in astrocyte-mediated synapse development, we discovered the Leucine-rich-repeat (LRR) protein family member *Lrig1* is a transcriptional target of Shh signaling in astrocytes. Lrig1 has previously been shown to regulate EGF signaling and dendrite outgrowth during nervous system development (Alsina et al. 2016, Del Rio et al. 2013). Leucine-rich-repeat (LRR) protein families are highly expressed throughout the CNS, where a number of them function as synaptic organizers (de Wit and Ghosh 2014). Whether astrocyte-derived *Lrig1* (or other Shh dependent genes) could function to promote the formation of bipartite or tripartite synapses between specific subsets of neurons and glial cells has yet to be determined. In future studies, it will be important to resolve the individual or combinatorial contributions of specific genes or functional classes of genes to Shh-dependent formation of specific synaptic connections.

We found that Il33 undergoes two phases of expression: first a Shh-dependent, astrocyte-specific phase in the developing cortex and then a Shh-independent, oligodendrocyte-specific phase in the mature cortex (**Figure 4 and S5**). This finding highlights the importance of accounting for the cell type and developmental stage specific expression patterns of Shh target genes when interrogating their function. Oligodendrocyte-specific Il33 has been shown to be required for oligodendrocyte maturation (Sung et al. 2019), while its expression in astrocytes was shown to regulate microglial synapse pruning (Vainchtein et al. 2018). However, our data suggests that activating Shh signaling in astrocytes produces a net increase in glutamatergic synapses. One potential mechanism that could reconcile our findings with the previously described function of Il33, is if specific types of synapses are repressed while the formation of other connections are facilitated. This differential impact on specific types of synapses might explain why ShhcKO leads to reduced VGluT2 synapses, but no change in VGluT1 (**Figure S7B and S7C**). In future studies it will be important to distinguish the inputs of intracortical neurons based upon their cortical areal location and molecular identity.

Astrocyte processes make intimate contacts with neuronal synapses where they are thought to both maintain homeostasis and modulate synaptic transmission. Our data shows that Shh signaling controls both the layer specific gross morphological complexity of cortical astrocytes and coverage of synapses by their fine processes. Not all cortical synapses are ensheathed by astrocytes; and the mechanisms that determine the types of synapses and the extent of synapse ensheathment are not known. One possibility is that Shh may be required for perisynaptic astrocyte process coverage of specific types of synapses. It has been reported that perisynaptic astrocyte processes are viewed as motile structures displaying rapid actin-dependent movements in response to neuronal activity (Bernardinelli et al. 2014, Reichenbach, Derouiche and Kirchhoff 2010, Haber, Zhou and Murai 2006). Thus, Shh may specifically influence the structure of perisynaptic astrocyte processes in an activity-dependent manner. However, in our data, we found that the fraction of synaptic interface perimeter surrounded by perisynaptic astroglia is reduced in Shh mutants, suggesting that less complex astrocytes might lead to reduced connections with surrounding neurons. This reduction in contact might influence the ability of astrocytes to perform homeostatic functions during synaptic transmission, such as rapid uptake of neurotransmitter (Perea, Navarrete and Araque 2009, Farhy-Tselnicker and Allen 2018).

During embryonic development, SHH is thought to disperse from floor plate cells to form a gradient that specifies cell types along the dorso-ventral axis of the developing CNS (Dessaud, McMahon and Briscoe 2008). It is not known whether similar mechanisms are at play to release SHH from the morphologically complex and polarized neurons in the cortex. Research on the developing limb bud suggests that SHH may be trafficked over long distances in cytonemes (Sanders, Llagostera and Barna 2013). A similar trafficking mechanism in neurons would provide an intriguing means for controlling the delivery of SHH to local astrocytes. Many Shh target genes including Gli1, Kir4.1, Il33 and Lrig1 have sharp boundaries of expression that demarcate neuronal layers Va and Vb (**Figure 3 and 4**). Our data is consistent with others, suggesting that least some portion of SHH protein is released by the somatodentritic compartments of cortical projection neurons (Rivell et al. 2019). When Shh was either deleted or overexpressed, the most robust gene expression changes were observed in the astrocytes located in close proximity to neuron cell bodies (**Figure 4 and 5**). Future studies focused on understanding the mechanisms that regulate the trafficking, sequestration and release of SHH protein in neurons will provide important insights into mechanisms of how neurons communicate with their astrocyte neighbors.

Our data shows that cortical neuron-derived Shh is developmentally regulated, with peak expression between the first and second postnatal weeks, after which it is substantially downregulated to the lower basal level found in adults (**Figure S1A**). Peak expression of Shh coincides with the peak period of cortical synaptogenesis in mouse (Farhy-Tselnicker and Allen 2018). However, the Shh receptors Ptch1 and Boc both maintain a high level of expression throughout adulthood (**Figure 1**). Boc remains exclusively expressed in neurons throughout development and adulthood (Harwell et al. 2012), while Ptch1 is initially exclusive to cortical astrocytes lineages and then begins to be expressed in adult layer IV neurons (**Figure S1C**). The functional relevance of Ptch1 expression in adult cortical neurons has yet to be explored. One intriguing possibility is that cell-type- and layer-specific responses to Shh signaling in the cortex may be further diversified by combinatorial expression of Shh receptors and interacting proteins. It is possible that Shh signaling in neurons and astrocytes during development is required to establish cortical circuitry, while signaling in the adult cortex may have roles in synaptic maintenance or plasticity. The respective neuron- and astrocyte-specific expression of Boc and Ptch1 during cortical development suggests a model whereby layer Vb neuron-derived Shh signals to intracortical projection neurons through Boc and to astrocytes through Ptch1. Evidence from previous research has shown that different downstream signaling pathways are activated through these receptors, with Ptch1 being required for transcriptional regulation of Gli1, while Boc is necessary for Src family kinase signaling (Yam et al. 2009). There are also instances in which Boc and Ptch1 have been shown to function cooperatively in *cis* (Allen et al. 2011, Izzi et al. 2011). Future studies are necessary to determine if *trans* interactions between these receptors allow Shh to function as a layer-specific organizer that coordinates both neurons and astrocytes to facilitate synaptic development. Taken together, our study highlights the critical role of neuron-derived Shh in regulating layer-specific astrocyte programs to coordinate neuron-glia communication during cortical circuit development.

## ACKNOWLEDGMENTS

The authors would like to thank Mahmoud El-Rifai and Aurélien Begué at the Neurobiology Imaging Facility at Harvard Medical School for the RNAscope experiments, Jasper Maniates-Selvin, Logan Thomos for electron microscopy technical support, Michelle Lowe Ocana at Neurobiology Imaging Facility for confocal imaging technical support. Clarence Yapp at Therapeutic Science Department at Harvard Medical School for MATLAB code and all members of the Harwell lab. Y.X. is partially supported by Alice and Joseph Brooks Fund Fellowship and Louis Perry Jones Postdoctoral Fellowship. Research in C.H.’s laboratory is supported by NIH Grants R01MH119156 and R01NS102228.

## AUTHOR CONTRIBUTIONS

Conceptualization, Y.X. and C.C.H.; Investigation, Y.X., A.K., W.W., Z.H., O.M., B.O., M.A., S.V., N.G., D.T, C.C., I.S.; Resources, M.T., W.A.L., B.S. and C.C.H.; Writing – original draft, Y.X. and C.C.H.; Writing – review & editing, Y.X., W.W., A.T.K., M.K., W.A.L., N.G., O.M., B.O., T.L., C.R., M.T.G., C.C.H.; Funding Acquisition, C.C.H.; Supervision, C.C.H.

## KEY RESOURCES TABLE

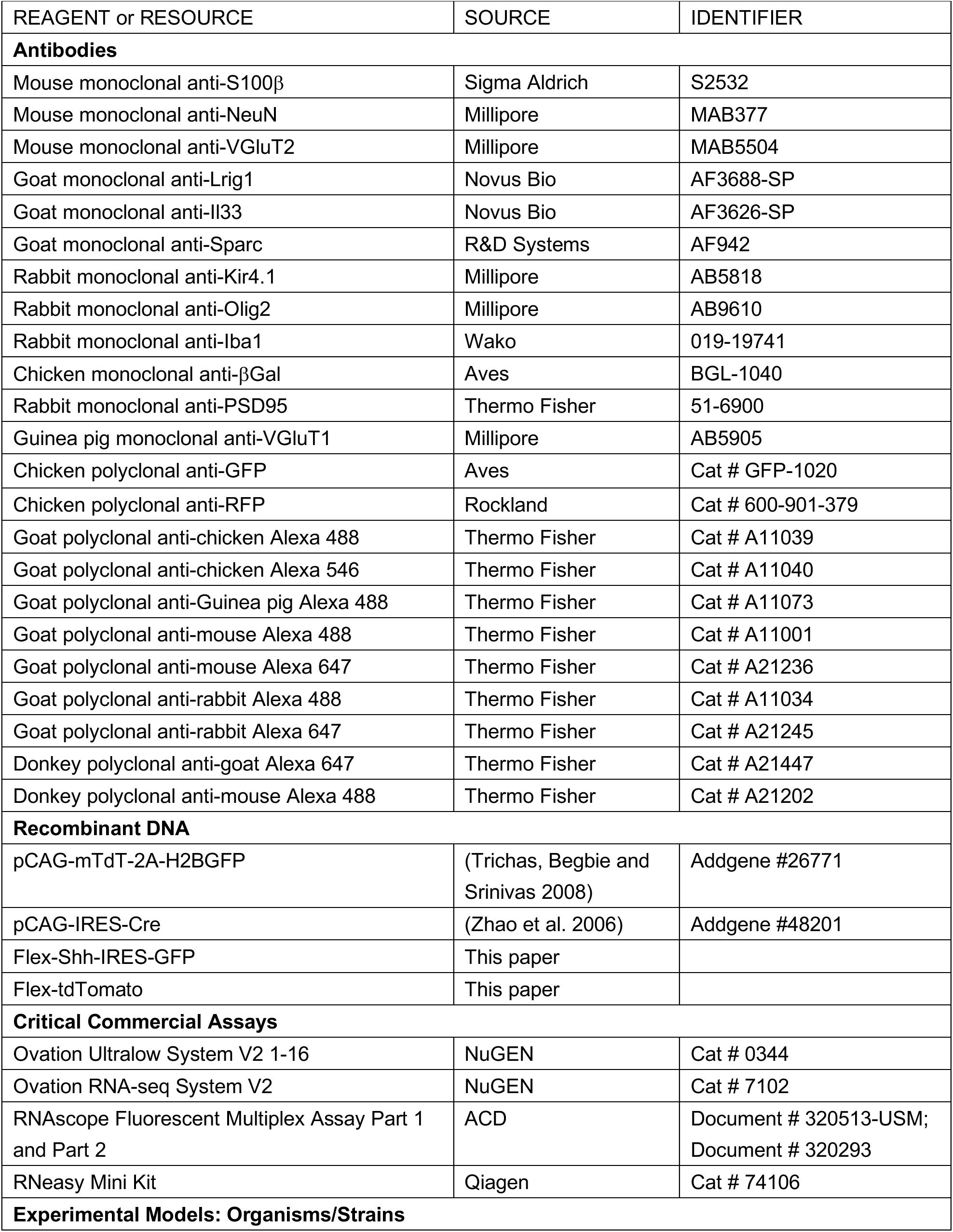
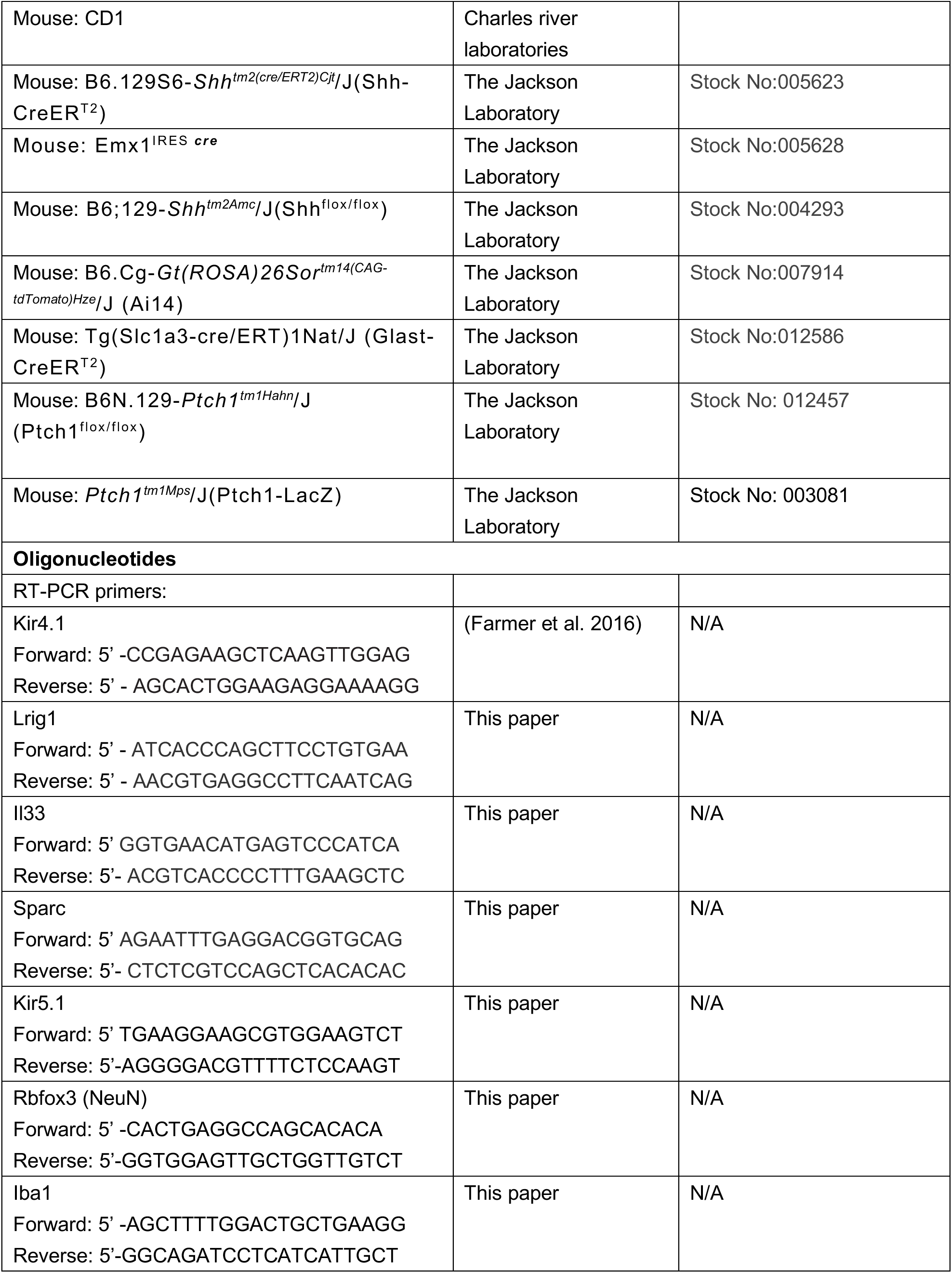
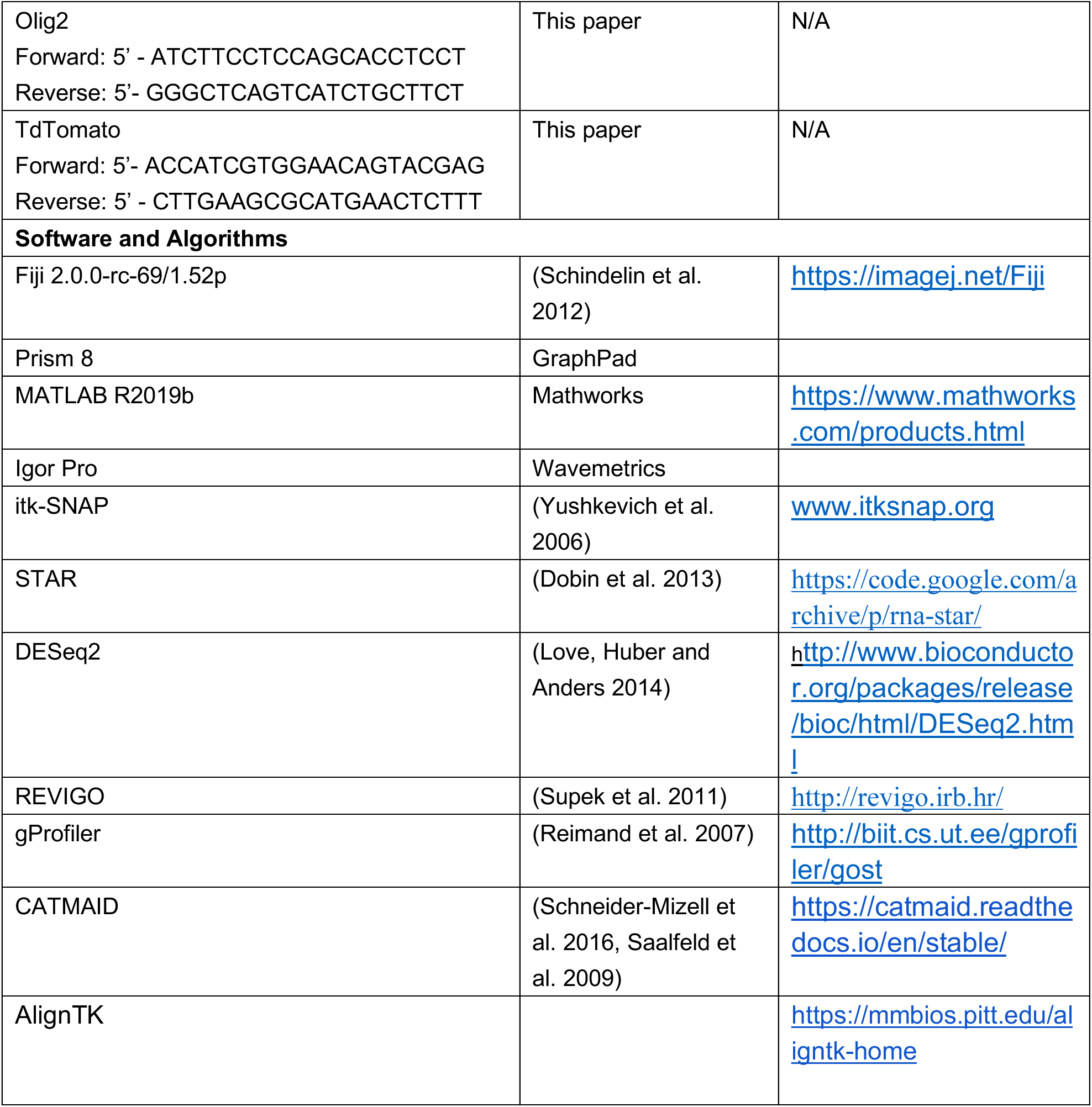

## LEAD CONTACT AND MATERRIALS AVAILABLILITY

This study did not generate new unique reagents. Further information and requests for resources and reagents should be directed to and will be fulfilled by the Lead Contact, Corey Harwell.

## EXPERIMENTAL MODEL AND SUBJECT DETAILS

All animal procedures conducted in this study followed experimental protocols approved by the Institutional Animal Care and Use Committee of Harvard Medical School. Mouse lines are listed in the Key Resources Table. Mouse housing and husbandry conditions were performed in accordance with the standards of the Harvard Medical School Center of Comparative Medicine. Mice were kept under temperature-controlled conditions on a 12:12 h light/dark cycle with food and water *ad libitum*. Samples were obtained from animals at postnatal days as indicated in Figure legends. In vivo experiments were performed on a batch of mice of either sex.

## METHOD DETAILS

### Tamoxifen administration

Tamoxifen (Sigma) was dissolved in corn oil at a concentration of 20 mg/ml at 37°C and stored at 4°C for the duration of injections. For P12 mice (WT : Glast-CreER^T2^; Ai14 or Ptch1cKO: Glast-CreER^T2^; Ptch1^flox/flox^; Ai14), 10-15 μL of tamoxifen was injected intraperitoneally for three consecutive days in order to induce effective recombination. For Shh-CreER^T2^; Ai14 mice, the dose of tamoxifen varied with mouse weight (age). Mice were closely monitored for any adverse reactions to the treatment.

### Immunofluorescence staining

Postnatal animals were transcardially perfused with PBS followed by 4% paraformaldehyde (PFA), their brains were dissected out and post-fixed in 4% PFA overnight at 4°C. Brains were sliced into 50 μm sections on a vibratome (Leica Microsystems VT1200S). Sections were prepared and placed in blocking solution (0.3% Triton (Amresco), and 10% goat serum) for 1-2 h at room temperature. After washing with PBS, sections were incubated sequentially with primary antibodies overnight at 4°C and secondary antibodies for 1 h at room temperature (RT). DAPI (4’,6-diamidino-phenylindole, Invitrogen) was added to the secondary antibodies solution. The sections were mounted using ProLong Gold Antifade Mountant (Invitrogen). Cryosections: Brains were cryoprotected in 30% sucrose/PBS overnight at 4°C after post-fixation. Brains were embedded in O.C.T. compound (Sakura), frozen on dry ice and stored at −80°C. Samples were sectioned at 20 μm on a cryostat (ThermoFisher CryoStart NX70). Images were acquired using a Leica SP8 laser point scanning confocal microscope,10X, 20X, 40X, 63X and 100X objectives were used and images were further analyzed using Fiji, brightness and contrast were adjusted as necessary for visualization, but the source images were kept unmodified.

### *In situ* hybridization (RNAscope)

Fresh frozen brain sections were prepared and pretreated by using the RNAscope fluorescent multiplex assay. RNA probes (see **Key Resources Table**) against *Lrig1, Kir5.1* were combined with a probe against an astrocyte marker *Aldh1l1*, for Ptch1cKO samples, multiplex RNAscope was done by combining *Lrig1 and Kir5.*1 probes with a *tdTomato* probe to confirm that labeled cells are recombined cells.

### Electron microscopy

Sample preparation: mice were transcardially perfused with Ames’ medium (oxygenated with 95% O_2_, 5% CO_2_, warmed to 37°C) to remove blood, and then a fixative solution (2.5% glutaraldehyde (Electron Microscopy Sciences) and 2% paraformaldehyde (Sigma) in cacodylate buffer (0.1 M sodium cacodylate (Electron Microscopy Sciences), pH 7.4 and 0.04% CaCl_2_ (Sigma), warmed to 37°C). Brains were dissected out and then post-fixed in the same fixative at 4°C overnight. After washing the samples with cacodylate buffer, brains were sectioned to 200 μm thickness using a vibratome, then stored in 3% glutaraldehyde in cacodylate buffer at 4°C before heavy metal staining. Heavy metal staining: sections were washed with cacodylate buffer. Then trimmed to include on the regions of interest (somatosensory cortex). Samples were then stained in 1% osmium tetroxide aqueous solution (Electron Microscopy Sciences) with 2.5% potassium ferrocyanide (MilliporeSigma) at RT for 1h. Sections were rinsed in water and then maleate buffer (MilliporeSigma) (pH=6.0) and stained in 0.05 M Maleate Buffer containing 1% uranyl acetate (Electron Microscopy Sciences) at 4°C overnight. Embedding: Sections were dehydrated in series of washes from 5% to 100% ethanol, then infiltrated with 1:1 Epon resin: propylene oxide at 4°C for overnight. Finally, sections were embedded in 100% Epon resin and polymerized at 60°C for 48–72 h. Serial sectioning and EM imaging: serial ultrathin sections (40-50 nm thick, 100-250 sections) were collected and imaged using the automated TEMCA-GT pipeline (Maniates-Selvin et al. 2020). Images were collected at 2500x magnification resulting in 4.3 nm pixels. Images were elastically aligned to form a 3D image volume using AlignTK software (see **Key Resources Table**) run on the O2 computing cluster at Harvard Medical School. Image analysis: Image volumes were accessed and annotated using the CATMAID software (see **Key Resources Table**). All synapses and astrocytes within regions of interest within each layer and sample (20×10×0.9 μm, axes correspond to medial-lateral, dorsal-ventral, and anterior-posterior directions, respectively) were annotated. For each synapse, the area of the post-synaptic density (PSD) was traced. Synapses whose cleft was surrounded by astrocytic processes were also labeled as astroglia synapses. For 3D visualization of synapses, image data surrounding the synapse was extracted using the pymaid API (https://pymaid.readthedocs.io/en/latest/), then manually segmented using itk-SNAP (see **Key Resources Table**). Only a few representative astroglia synapses were reconstructed this way. Astrocyte contact fraction analysis: astrocytic perimeter was measured by summing the length of the astrocyte contacts at the edges of synaptic axon-dendrite interface. And the total perimeter was measured by summing the astrocyte contact length and the edges of axon-dendrite, then the fraction of astrocytic contact was determined. Glycogen granules analysis: a single cross-section of each tripartite synapse with a prominent PSD was manually segmented using itk-SNAP. Astrocytic glycogen granules were identified as dark, round spots, 50-80 nm in diameter, that were distributed over a relatively clear cytoplasm.

### RNA-Seq

Brains were dissected out after transcardial perfusion of animals with PBS and sliced into coronal sections with a 1.0 mm brains slicer Matrix (BSMAS001-1). A 1 mm X 2 mm piece of somatosensory cortex was cut off and minced tissues into small pieces. Cortical tissues were digested with accutase (ThermoFisher) on a rotator at 4°C. Cells were centifuged at 2000 rpm for 2 min at 4°C. The cell pellet was resuspended in ice cold Hibernate-A (ThermoFisher) and triturated with a 1 ml pipette tip until the cortical chunks disappeared. The cell suspension was gently pipetted out through a 70 μm filter, and ∼800K cells were sorted by tdTomato fluorescence, tdTomato– cells were sorted out as well to use as negative controls. RNA was then purified with a RNeasy Mini Kit (Qiagen). The concentration and quality of purified RNA were determined using BioAnalyzer (Agilent), and the RNA was reverse-transcribed into cDNA and amplified by RNA-based single primer isothermal amplification (SPIA) using the Ovation RNA-seq system V2 (NuGEN). Synthesized cDNA was sonicated using a Covaris S2 ultrasonicator to reduce the fragment size range to 100-600 bp. 100 ng of sheared cDNA were end paired, ligated with barcoded adaptors, amplified and purified using the Ovation Ultralow System V2 (NuGEN). RNA Exome Capture and Sequencing: targeted exome hybrid capture was performed using Illumina TruSeq Exome Capture reagents according to manufacturer’s protocol on a Beckman Coulter Biomek FX. Post capture the library pools were quantified by Qubit fluorometer, Agilent TapeStation 2200, and RT-qPCR using the Kapa Biosystems library quantification kit according to manufacturer’s protocols. Uniquely indexed libraries were pooled in equimolar ratios and sequenced on an Illumina NextSeq 500 with single-end 75bp reads by the Dana-Farber Cancer Institute Molecular Biology Core Facilities. Libraries were sequenced in an Illumina NextSeq 500 to a sequencing depth of 28-40 million reads per sample.

### RNA-seq Analysis

Sequenced reads were aligned to the UCSC mm10 reference genome assembly and gene counts were quantified using STAR (v2.5.1b) (Dobin et al. 2013). Differential gene expression testing was performed by DESeq2 (v1.10.1) (Love et al. 2014) and normalized read counts (FPKM) were calculated using cufflinks (v2.2.1) (Trapnell et al. 2010). RNAseq analysis was performed using the VIPER snakemake pipeline (Cornwell et al. 2018). Significant genes were identified using a threshold of P<0.05, Lists of differentially expressed genes were examined by gProfiler for statistical enrichment of information such as Gene Ontology (GO) terms, biological pathways functional redundancy in over-represented GO terms identified by gProfiler was reduced with REVIGO for a visual representation of the most prominent processes.

### Quantitative reverse transcriptase polymerase chain reaction (qRT-PCR)

The somatosensory cortex of WT and ShhcKO mice was dissected out in PBS and stored at −80°C. Total RNA was extracted using the Qiagen RNeasy Mini Kit. First strand synthesis was performed using Invitrogen a SuperScript III First-Strand Synthesis Kit with Oligo(dT)_20_ primers (Invitrogen). Quantitative PCR was performed using Sybr Green Supermix (BIO-RAD) on a C1000 Touch thermocycler (BIO-RAD). Relative levels of mRNA were calculated using the delta CT method with *Gapdh* as the internal control. For RNA-seq samples, a subset of genes was confirmed by real-time PCR analysis using purified cDNA of sorted cells, Primers designed for each tested gene can be found in the **Key Resources Table**.

### *In utero* electroporation

Timed pregnant mice were anesthetized using an isoflurane vaporizer and placed on a warming pad. An abdominal incision of about 1 inch in length was made and the uterine horns were carefully exposed on top of a sterile gauze pad. Embryos were kept moist with pre-warmed PBS at 37°C during the entire procedure. Approximately1-2 μL of endotoxin-free DNA (1-3 mg/ml) diluted in PBS/0.025% Fast Green (SIGMA) was injected into the lateral ventricles of the forebrain using heat-pulled glass micropipettes (Drummond). Once all embryos were injected, 5 pulses of 30-40 V (50 ms duration and 950 ms intervals) were applied with 7 mm platinum electrodes (BTX) connected to an ECM 830 square wave electroporator (BTX). The abdominal cavity was then sutured and stapled before administering buprenorphine (0.05-0.1 mg/kg) and ketoprofen (5-10 mg/kg). Mice were allowed to recover in a 37°C chamber after surgery. All mice were collected at P14.

### Postnatal astrocytes label by electroporation (PALE)

Early postnatal (P0-P1) WT (Shh^flox/flox^) and ShhcKO (Emx1-Cre; Shh^flox/flox^) mice were sedated by hypothermia by being placed on ice for 4-5 min. Approximately 1 μL of endotoxin-free plasmid DNA (1 mg/ml) with Fast Green Dye was injected into the lateral ventricles of one hemisphere as described above (*In utero* Electroporation method). We used a plasmid (CAG-mTdT-2A-H2BGFP) in order to label nuclei with GFP and astrocyte processes with tdTomato. Following the plasmid injection, 5 discrete 50 msec pulses of 100 V were delivered at 950 msec intervals to the head of each injected pup, with the positive pole of the electrode placed on the side of the injection. Pups were covered on a 37°C pad until fully responsive, then placed back into their home cage.

### In-vivo astrocyte complexity analysis (Sholl analysis)

High magnification 63X objective Z-stack images were obtained using a Leica SP8 confocal microscope. 5 μm-thick Z-stacks of 18 serial sections were imaged with tdTomato and GFP (WT and ShhcKO) or tdTomato (WT and Ptch1cKO) channels. Each Z-stack was converted into maximum projection images using FIJI. Sholl analysis was performed using the Sholl analysis plugin in FIJI. Starting sholl radii at 5 μm from the line start, sholl radii at 1μm increments.

### Analysis of synaptic staining

For electroporated mice, P21 controls and ShhcKO sections were stained with an antibody against pre- and postsynaptic makers: VGluT1 and PSD95, VGluT2 and PSD95. High magnification (100 × 1.44 NA objective plus 1.5X optical zoom) Z-stack images were obtained with a Leica SP8 confocal microscope. Each image including individual astrocyte with 5 um-thick Z stacks. The number of co-localized synaptic puncta on top of tdTomato+ astrocyte territory was analyzed using MATLAB (written by Yapp Clarence). For tamoxifen induced mice (WT and Ptch1cKO), high magnification of 2D images were obtained by a Leica Sp8 confocal microscope, the coding applies to either 2D or 3D images.

### Electrophysiology

#### Acute Brain Slice Preparation

Brain slices were obtained from P21-P28 days old mice, WT and Ptch1cKO (both male and female) using standard techniques. Mice were anesthetized by isoflurane inhalation and perfused transcardially with ice-cold artificial cerebrospinal fluid (ACSF) containing (in mM) 125 NaCl, 2.5 KCl, 25 NaHCO_3_, 2 CaCl_2_, 1 MgCl_2_, 1.25 NaH_2_PO_4_ and 25 glucose (295 mOsm/kg). Brains were blocked and transferred into a slicing chamber containing ice-cold ACSF. Coronal slices of cerebral cortex were cut at 300 μm thickness with a Leica VT1000s vibratome in ice-cold ACSF, transferred for 10 min to a holding chamber containing choline-based solution (consisting of (in mM): 110 choline chloride, 25 NaHCO_3_, 2.5 KCl, 7 MgCl_2_, 0.5 CaCl_2_, 1.25 NaH_2_PO_4_, 25 glucose, 11.6 ascorbic acid, and 3.1 pyruvic acid) at 34C then transferred to a secondary holding chamber containing ACSF at 34°C for 10 min and subsequently maintained at room temperature (20–22°C) until use. Electrophysiology recordings: Individual brain slices were transferred into a recording chamber, mounted on an upright microscope (Olympus BX51WI) and continuously superfused (2–3 ml min^−1^) with ACSF warmed to 32–34°C by passing it through a feedback-controlled in-line heater (SH-27B; Warner Instruments). Cells were visualized through a 60X water-immersion objective with either infrared differential interference contrast optics or epifluorescence to identify tdTomato+ cells. For whole cell voltage clamp recording, patch pipettes (2–4 MΩ) pulled from borosilicate glass (G150F-3, Warner Instruments) were filled with internal solution containing (in mM) 135 CsMeSO_3_, 10 HEPES, 1 EGTA, 3.3 QX-314 (Cl− salt), 4 Mg-ATP, 0.3 Na-GTP, 8 Na_2_-phosphocreatine (pH 7.3 adjusted with CsOH; 295 mOsm·kg−1). mEPSCs were recorded for 5min in the presence of 1 μM tetrodotoxin (Tocris) and 10 μM gabazine (Tocris) at a holding potential of –70 mV. To record evoked EPSC of pyramidal neurons in Layer V of S1 region, the membrane voltages were clamped at –70 mV, and 10 μM gabazine were added to the bath, extracellular stimulation was performed with a stimulus isolation unit (MicroProbes, ISO-Flex), bipolar electrodes (100 μm apart, PlasticOne) were placed on layer II/III of S1, 200 to 400 μm perpendicularly away from recording cells, evoke EPSC input-out curves were generated at 0.1 ms, from 10 μA at 10 μA steps to 100 μA. Data acquisition and analysis: membrane currents and potentials were amplified and low-pass filtered at 3 kHz using a Multiclamp 700B amplifier (Molecular Devices), digitized at 10 kHz and acquired using National Instruments acquisition boards and a custom version of ScanImage written in MATLAB. Electrophysiology data were analyzed offline using Igor Pro. Voltage-clamp traces represent the averaged waveform of five to seven consecutive acquisitions (**Figure 7**). Detection threshold for mEPSCs was set at 7 pA. Averaged waveforms were used to obtain current latency and peak amplitude. Peak amplitudes were calculated by averaging over a 2 ms window around the peak. Grubb’s test was used to detect outliers in this data. Data were compared statistically by Welch’s t test.

## SUPPLEMENTARY FIGURE LEGENDS

**Figure S1:**
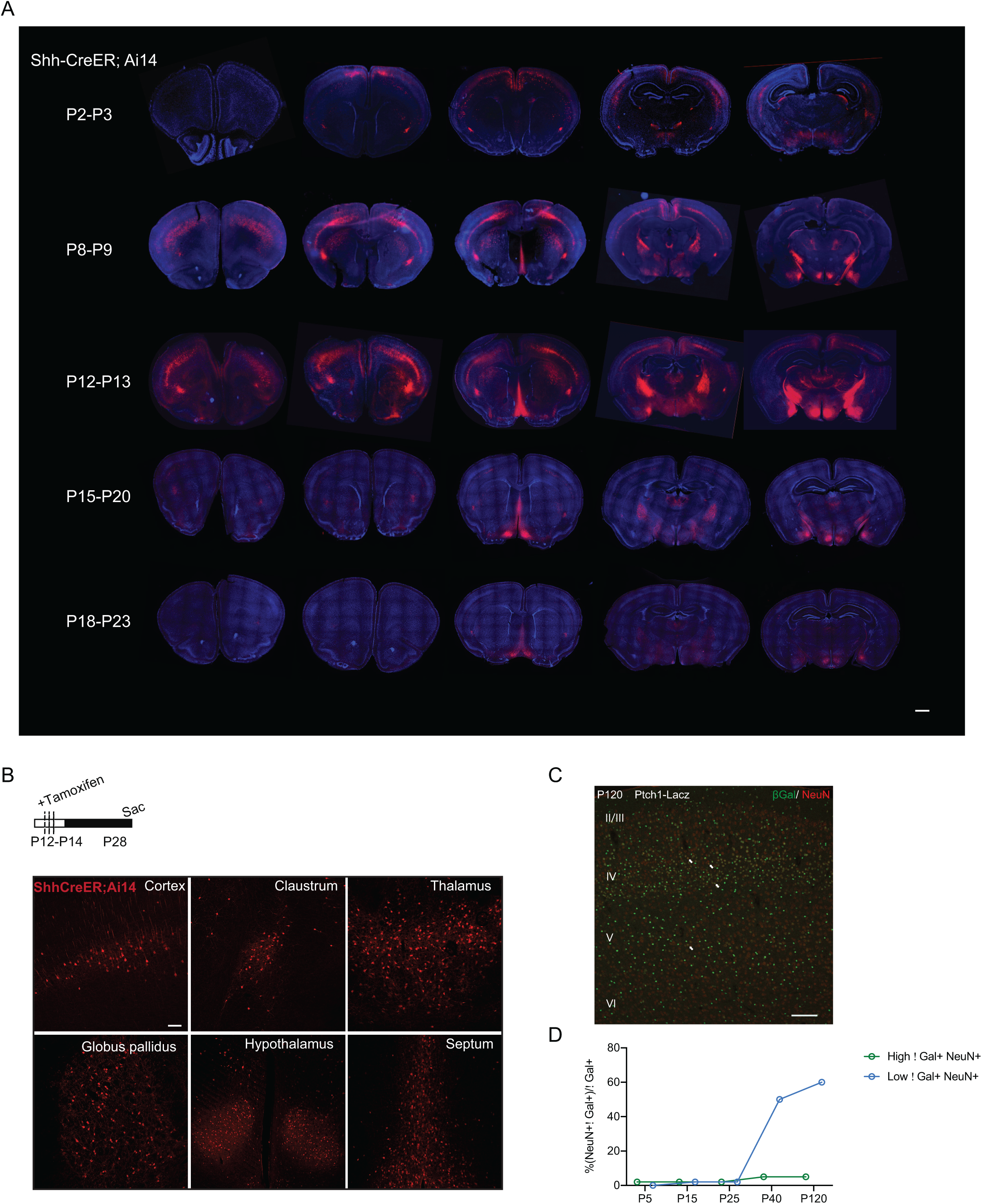
Spatial and temporal expression pattern of Sonic hedgehog in developing brains. **A**) Shh expressing cells are labelled with tdTomato fluorescence after injecting tamoxifen into mice (Shh-CreER^T2^; Ai14) at different age (P2-P3, P8-P9, P12-P13, P15-P20, P18-P23). Brains were collected at P28. Images were acquired using VS120 whole slide scanning. Scale bar: 1 mm. **B**) Shh expressing cells are labelled with tdTomato fluorescence after administrating tamoxifen at P12-P14 to efficiently induce Cre recombination. TdTomato+ cells are located in different brain regions, including layer V cortex, claustrum, thalamus, globus pallidus, hypothalamus and septum at P28 brains. Scale bar: 100 μm. **C**) Weakly labelled βGal+ (green) cells that are positive for NeuN (red) in P120 Ptch1-LacZ mice. Cortical layers are indicated in white. Scale bar: 250 μm. **D**) Quantification of percentage of NeuN+βGal+ cells in total βGal+ cells. High βGal+NeuN+ cells (P5: 2%, P15: 2%, P25: 2%, P40: 5%, P120: 5%) and low βGal+NeuN+ cells (P5: 0%, P15: 2%, P25: 2%, P40: 50%, P120: 60%).

**Figure S2:**
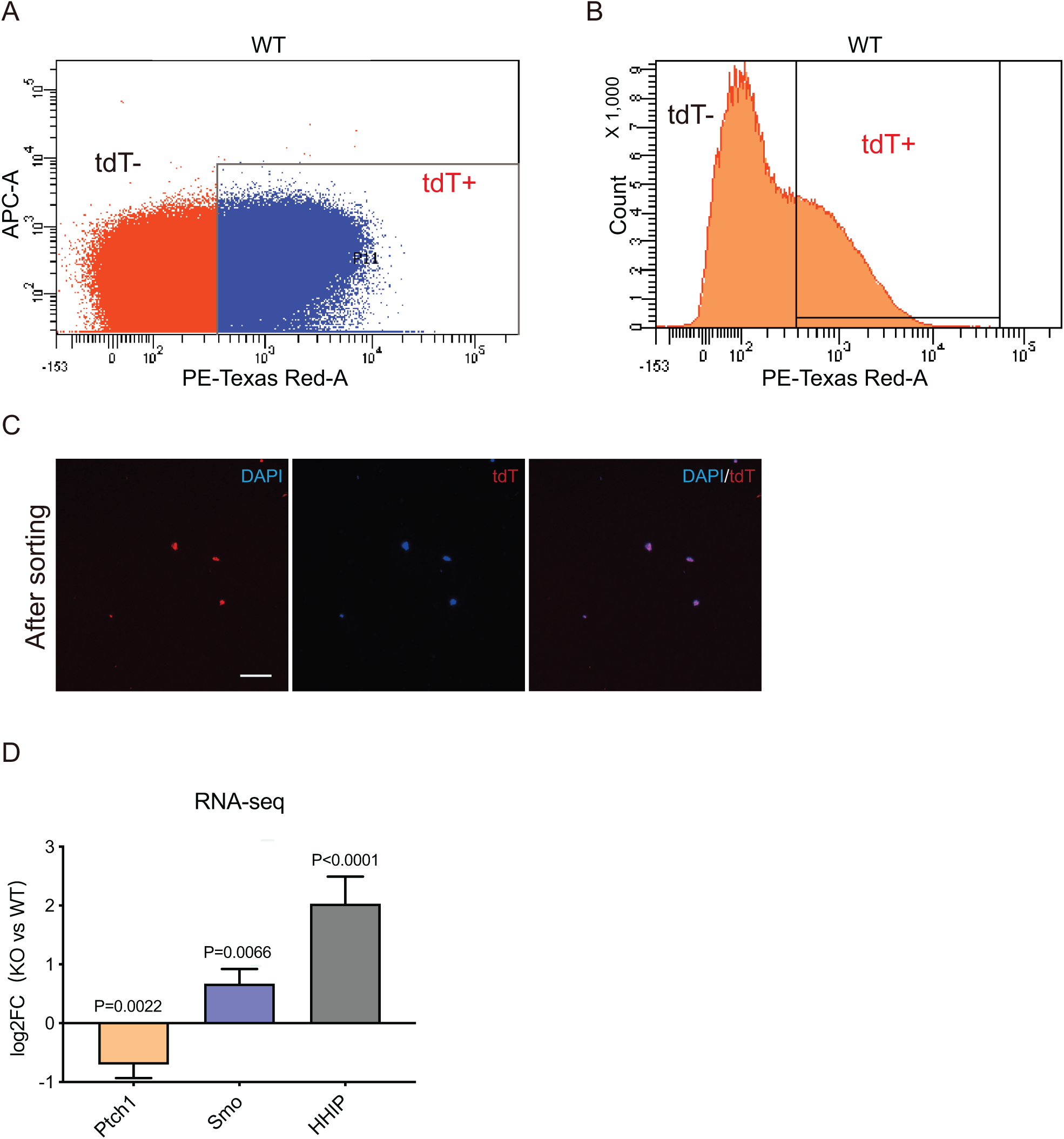
Transcriptional profiling of WT and Ptch1cKO astrocytes during cortical development. **A**, **B**) Representative plots showing sorting gates for tdTomato+ and tdTomato– cells. **C**) Representative images of cells suspensions stained for tdTomato (red) and DAPI (blue) after sorting. **D**) Fold-changes of *Ptch1*, *Smo* and *Hhip* expression in Ptch1cKO astrocytes.

**Figure S3:**
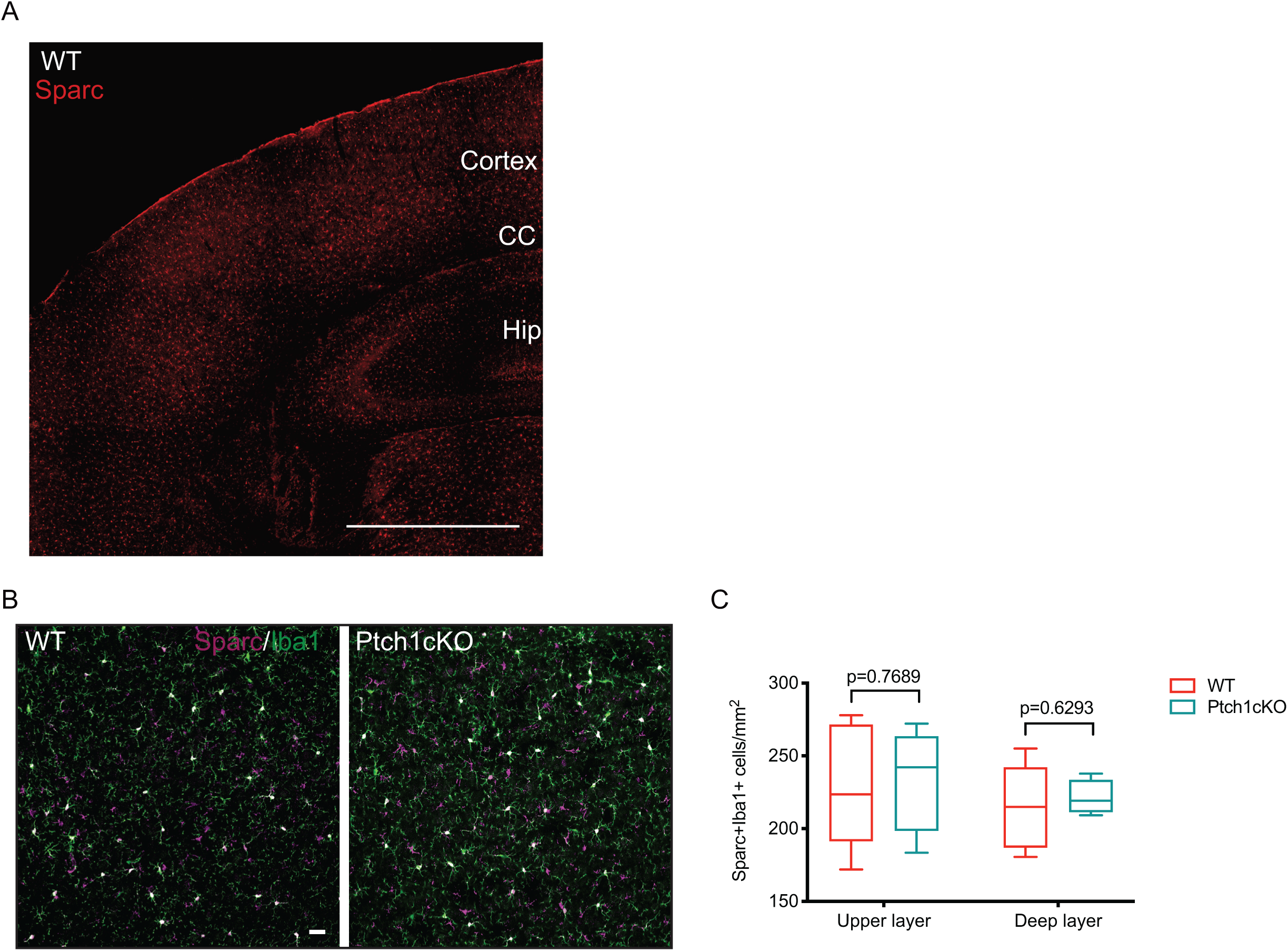
Sparc is highly expressed in microglia in the developing cortex. **A**) Immunostaining shows that Sparc (red) is highly expressed in P21 cortex through upper and deep layers. Scale bar: 1mm; **B**) Immunostaining shows that Sparc is enriched in Iba1+ microglia cells. **C**) Number of Sparc+Iba1+ cells is not changed in Ptch1cKO compared to WT. The mice were injected with tamoxifen at P12-P14. N=6 mice for each condition. Deep layer: WT (230.19 ± 20.11), KO (300.39 ± 17.84); upper layer: WT (195.81 ± 21.87), KO (265.05 ± 11.07). Data represent mean ± SEM; statistical analysis is multiple t-test.

**Figure S4:**
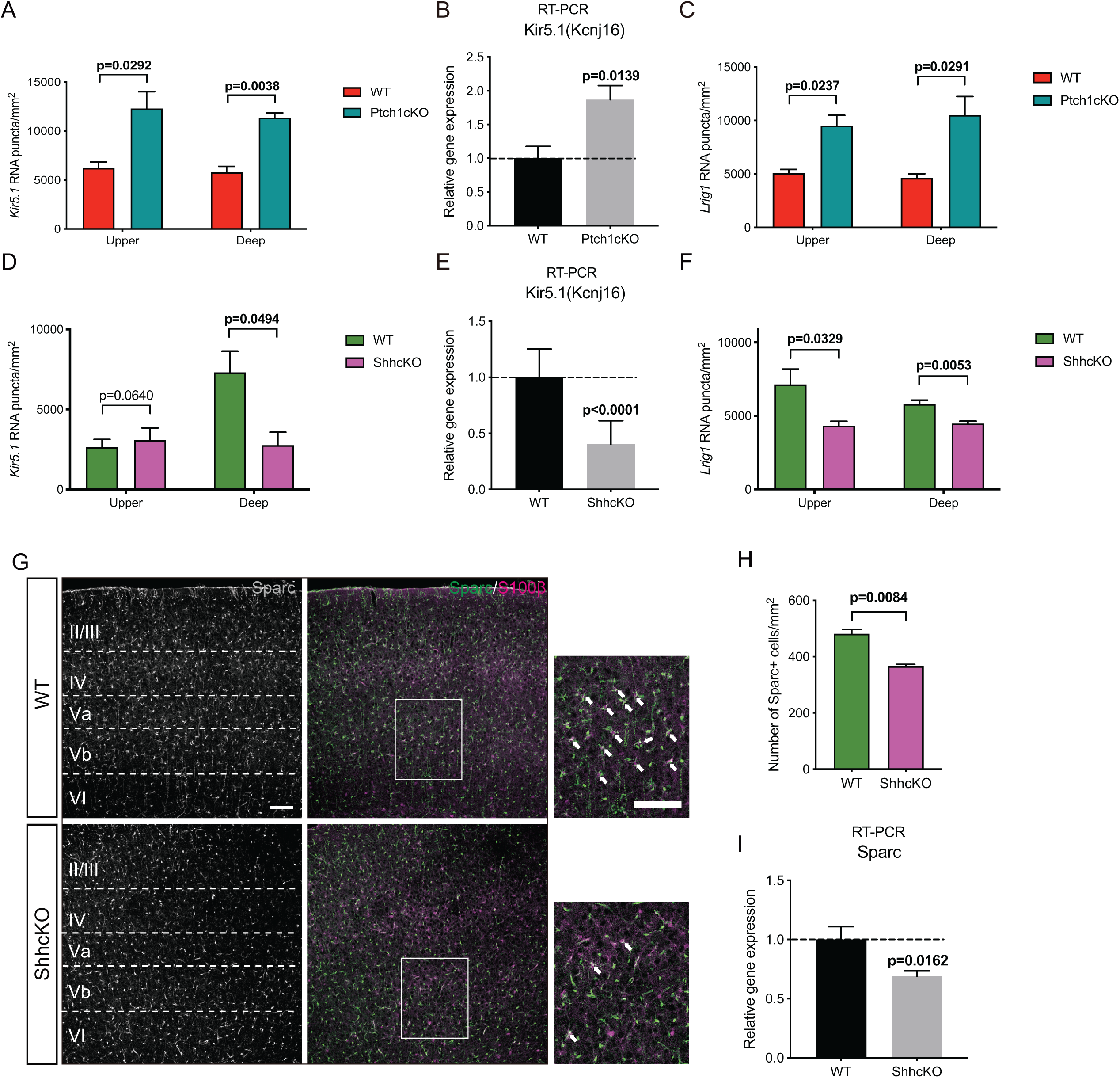
Shh regulates target genes expression in cortical astrocytes. **A**) In situ hybridization shows that the number of *Kir5.1* puncta per mm^2^ is increased in both upper and deep layer of P21 Ptch1cKO. The mice were injected with tamoxifen at P12-P14. N=3 mice for each condition. Deep: WT (5785.19 ± 606.29), KO (11370.37 ± 478.61); upper: WT (6237.04 ± 609.56), KO (12294.44 ± 1716.94). **B**) RT-PCR shows a significant increase in the level of *Kir5.1* RNA in P21 Ptch1cKO cortex. The mice were injected with tamoxifen at P12-P14. WT (N=4 mice, 1.00 ± 0.18), KO (N=4 mice, 1.87 ± 0.21). **C**) *Lrig1* RNA was evaluated in P21 Ptch1cKO using *in situ* hybridization. Number of *Lrig1* puncta per mm^2^ is increased in both upper and deep layers of P21 Ptch1cKO cortex. The mice were injected with tamoxifen at P12-P14. N=3 for each condition. Deep: WT (4631.48 ± 379.44), KO (10514.81 ± 1725.65); upper: WT (5076.54 ± 338.24), KO (9520.37 ± 957.87). **D**) Number of *Kir5.1* puncta per mm^2^ is reduced in deep layers of P15 ShhcKO cortex, while there is no significant change in upper layers. N=4 mice for each condition. Deep: WT (7312.04 ± 1297.20), KO (2763.89 ± 816.15); upper: WT (2634.26 ± 492.96), KO (3077.78 ± 755.73). **E**) There is a significant reduction in RNA level of *Kir5.1* in P15 ShhcKO cortex using RT-PCR. WT (N=4 mice, 1.00 ± 0. 25), KO (N=4 mice, 0.40 ± 0.21). **F**) Number of *Lrig1* puncta per mm^2^ is reduced in both upper and deep layer of P15 ShhcKO. N=5 mice for each condition. Deep: WT (5811.11 ± 262.21), KO (4478.89 ± 168.28); upper: WT (7133.70 ± 1046.20), KO (4328.15 ± 305.95). **G**) Immunostaining for Sparc (grey/green) and S100β (magenta) in WT and ShhcKO cortex. Cortical layers are indicated in white. High magnification images are shown in right panels, white arrows indicate Sparc+S100β+ cells, Scale bar: 100 μm. **H**) There is a significant reduction in the number of Sparc+ cells in ShhcKO cortex. WT (N=3 mice, 481.87 ± 15.03), KO (N=4 mice, 366.78 ± 6.11). **I**) RT-PCR shows a reduction in expression of *Sparc* in ShhcKO. WT (N=4 mice, 1.00 ± 0. 17), KO (N=4 mice, 0.59 ± 0.02). Data represent mean ± SEM; statistical analysis are multiple t-test or Welch’s t test.

**Figure S5:**
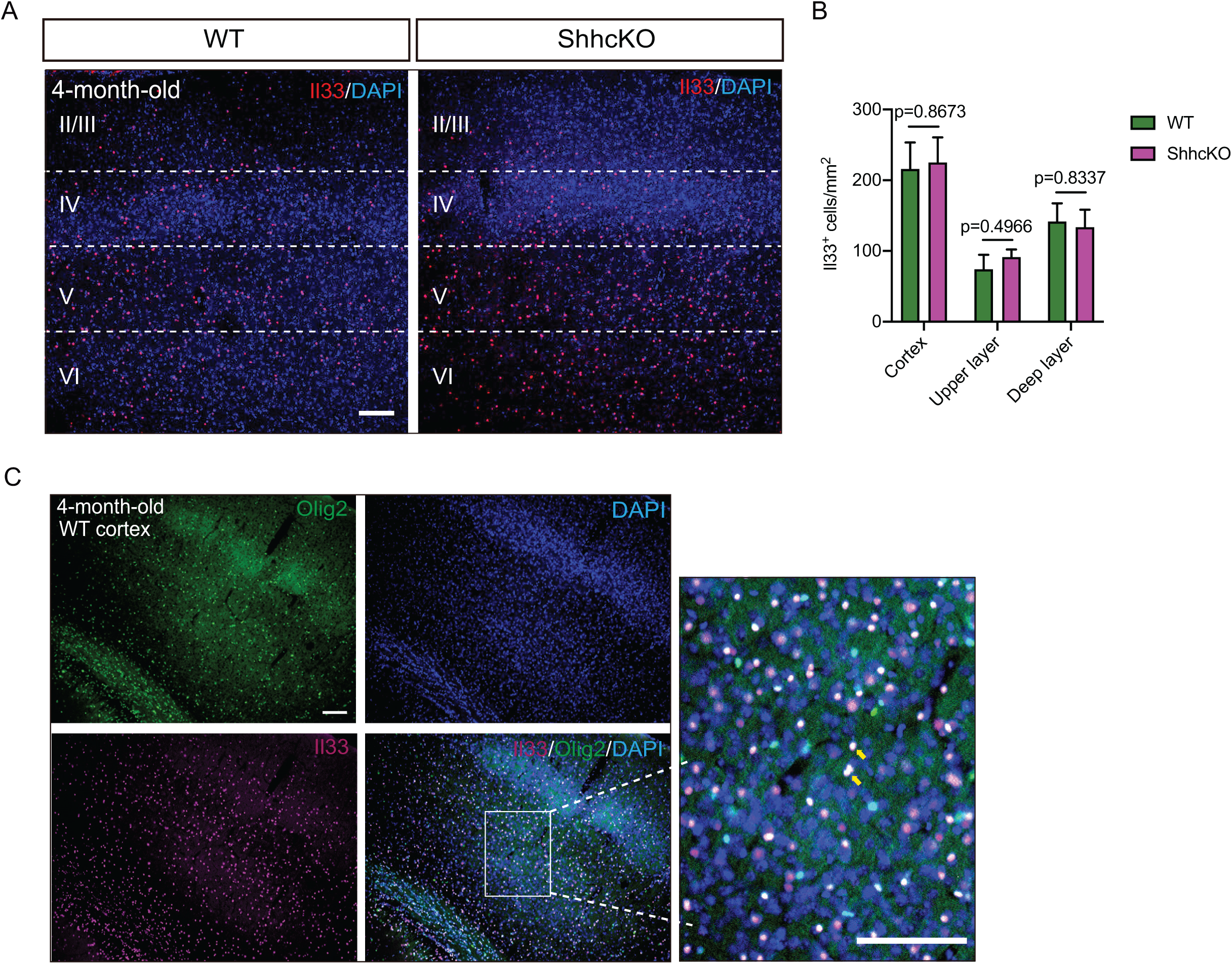
Il33 is highly expressed in oligodendrocytes in adult cortex. **A**) Immunostaining for Il33 (red) and DAPI (blue) in 4-month-old cortex from WT and ShhcKO mice. Il33 is highly enriched in adult cortex. Scale bar: 100 μm. **B**) Number of Il33+ cells per mm^2^ is not significantly different between ShhcKO and WT cortex in 4-month-old mice, either upper or deep layers. N=3 mice for each condition. Cortex: WT (215.92 ± 37.54), KO (225.11 ± 35.42); upper layer: WT (74.24 ± 20.30), KO (91.40 ± 10.78); deep layer (141.68 ± 25.64), KO (133.71 ± 24.69). **C**) Immunostaining for Il33 (magenta), Olig2 (green) and DAPI (blue) in 4-month-old WT, Il33 overlaps with Olig2, labelling oligodendrocytes across the cortex. High magnification images are shown in right panels. Yellow arrows indicate Il33+Olig2+ cells. Scale bar: 100 μm. Data represent mean ± SEM; statistical analysis is multiple t-test.

**Figure S6:**
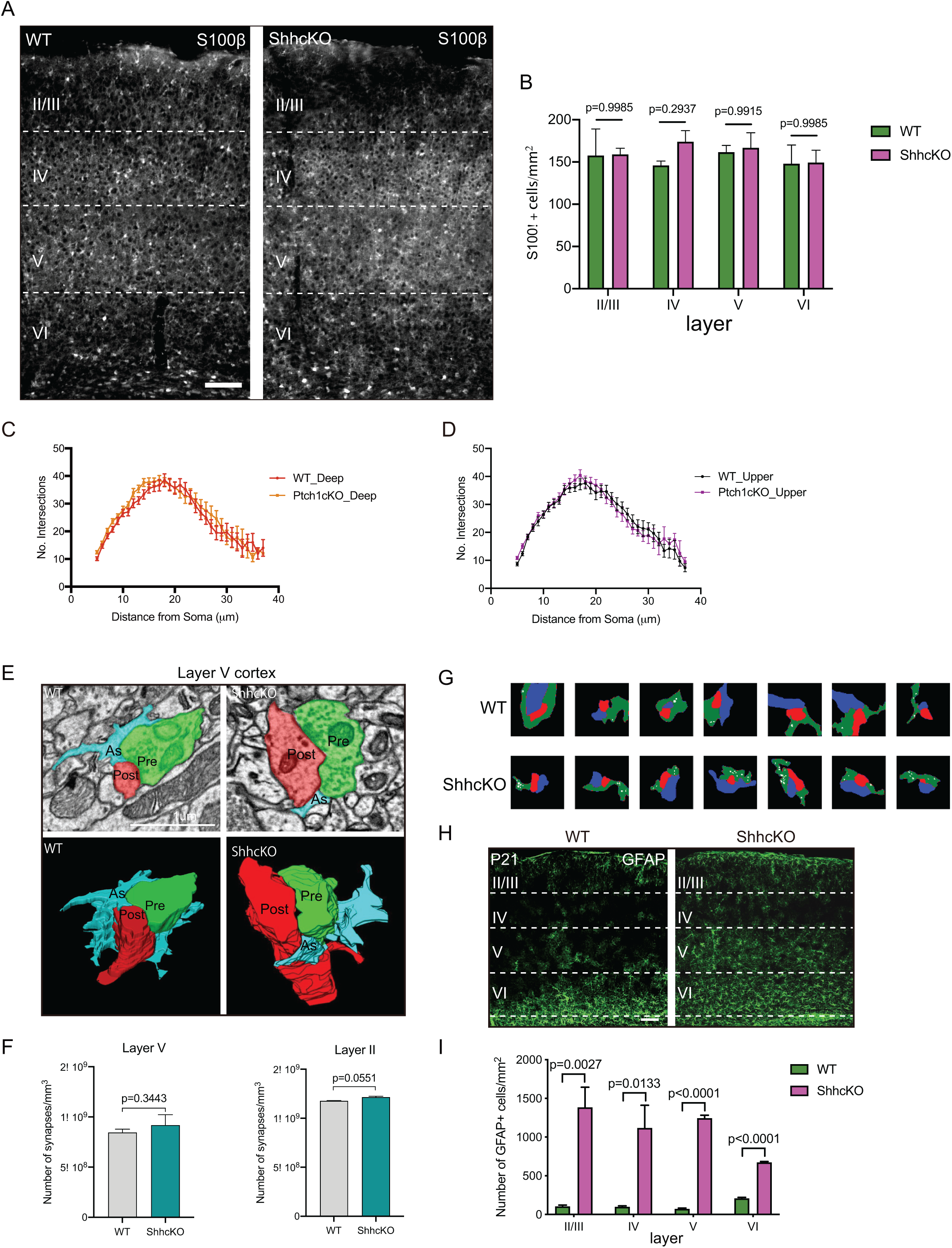
Deletion of Shh changes morphogenesis of astrocytes in developing cortex. **A**) Immunostaining for S100β (grey) in P21 WT and ShhcKO cortex. Cortical layers are indicated in white. Scale bar: 100 μm. **B**) Number of S100β+ cells per mm^2^ is not changed in ShhcKO through layers II/III to VI. N= 5 mice for each condition. Layer II/III: WT (157.65 ± 31.51), KO (158.95 ± 7.31); layer IV: WT (145.92 ± 5.19), KO (173.93 ± 13.17); layer V: WT (161.56 ± 8.02), KO (166.77 ± 17.76); layer VI: WT (147.88 ± 22.15), KO (149.18 ± 14.76). **C**, **D**) Sholl analysis of tdTomato labelled deep layer (**C**) and upper layer astrocytes (**D**) indicates there is no difference in astrocyte complexity between P21 WT and Ptch1cKO. The mice were injected with tamoxifen at P12-P14. Deep: WT (N=4 mice, n=32 cells), KO (N=4 mice, n=36 cells); upper: WT (N=4 mice, n=20 cells), KO (N=4 mice, n=23 cells). **E**) Representative 3D reconstruction of tripartite synapses from deep layers of WT and ShhcKO cortex. Astrocytes (blue) ensheath the nearby pre-(green) and post-synapses (red). Scale bar: 1 μm. **F**) Number of synapses per mm^3^ is not changed in either layer II or layer V of ShhcKO cortex compared to WT. N=1 mouse for each condition, n=3 boxes (20X10X0.9 μm) in layer II and layer V respectively. Layer V: WT (8.44 × 10^8^ ± 1.95 × 10^7^), KO (9.19 × 10^8^ ± 6.02 × 10^7^); Layer II: WT (1.18 × 10^9^ ± 3.34 × 10^6^), KO (1.22 × 10^9^ ± 1.03 × 10^7^). **G**) Representative 2D images of tripartite synapses from WT and ShhcKO. Presynaptic axons are in blue, postsynaptic dendrites are in red, astrocytes with white glycogen granules are in green. **H**) Immunostaining for GFAP in P21 WT and ShhcKO cortex, cortical layers are indicated in white. Scale bar: 100 μm. **I**) Number of GFAP+ astrocytes in WT and ShhcKO in indicated layers. N=4 mice for each condition. Layer II/III: WT (103.771 ± 17.08), KO (1384.18 ± 260.77); layer IV: WT (99.07 ± 12.04), KO (1118.14 ± 293.17); layer V: WT (72.20 ± 9.52), KO (1243.62 ± 38.92); layer VI: WT (207.82 ± 13.06), KO (672.11 ± 11.30). Data represent mean ± SEM; statistical analysis is multiple t-test (**B**, **C**, **D and I**) and Welch’s test (**F**).

**Figure S7:**
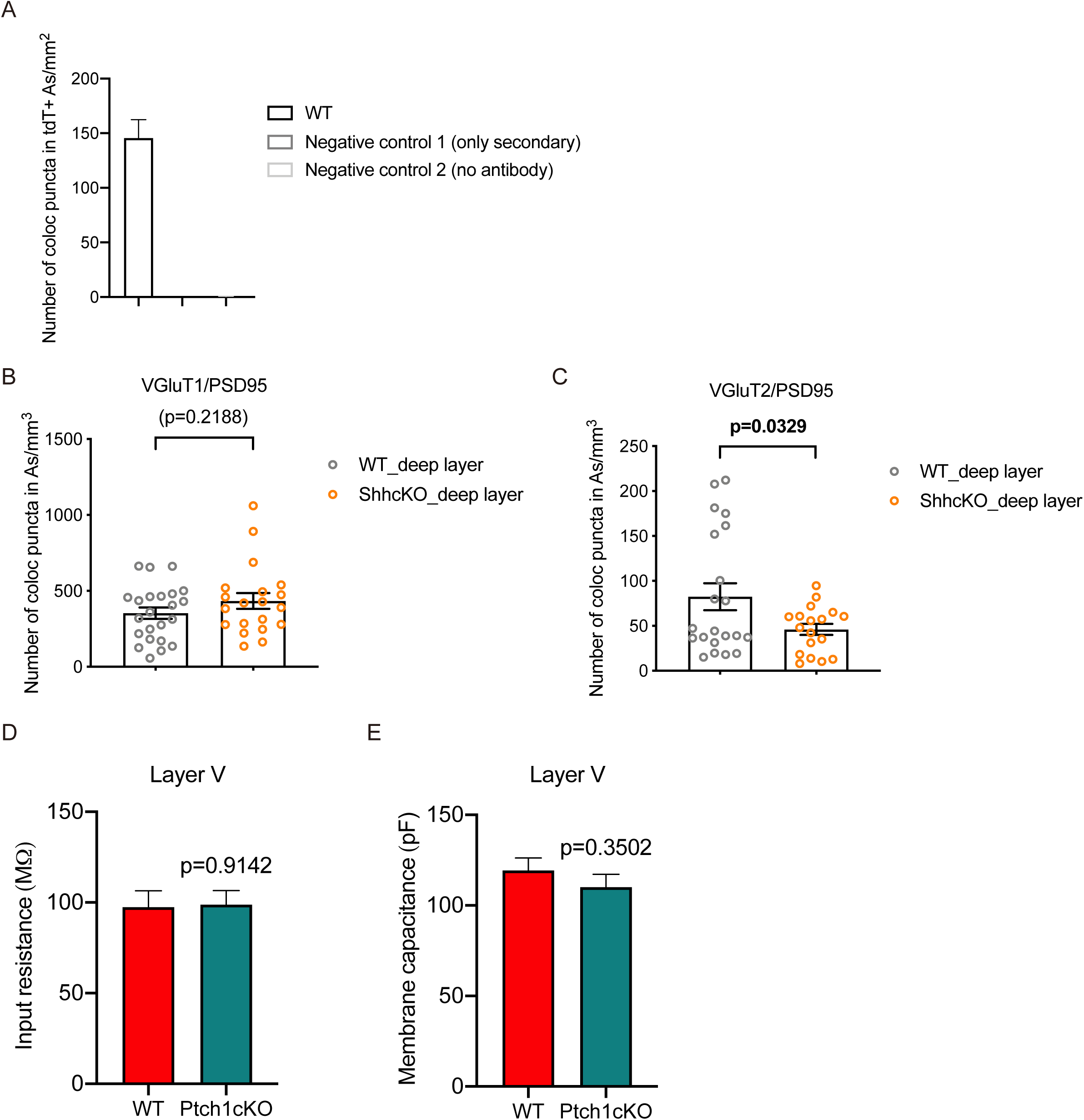
Activation of Sonic signaling in astrocytes promotes cortical synapse formation. **A**) The mice were injected with tamoxifen at P12-P14. Quantification of colocalized puncta (VGluT1/PSD95) in tdTomato+ astrocyte territory. Both negative control 1 (only secondary) and 2 (no antibody), it shows minimal amounts of false positives. N=3 mice for each condition. **B**) Number of VGluT1/PSD95 colocalized puncta in astrocyte domains per mm^3^ is not changed in deep layers of ShhcKO cortex. N=3 mice for each condition (WT: n=23 cells, 353.34 ± 37.73; KO: n=20 cells, 433.66 ± 51.91). **C**) Number of VGluT2/PSD95 colocalized puncta in astrocyte domains per mm^3^ is reduced in deep layers of ShhcKO cortex (WT: n=21 cells, 82.47 ± 14.98; KO: n=18 cells, 46.03 ± 6.14). **D**, **E**) Input resistance (**D**) and membrane capacitance (**E**) are not affected in layer V neurons of ShhcKO cortex. N=3 mice for each condition. Input resistance: WT, n=20 cells, 97.47± 8.98; KO: n=22 cells, 98.76 ± 7.76); membrane capacitance: WT (n=20 cells, 119.27 ± 6.84), KO (n=22 cells, 110.05 ± 7.09); Data represent mean ± SEM; statistical analysis is Welch’s t test.

**Table S1.**
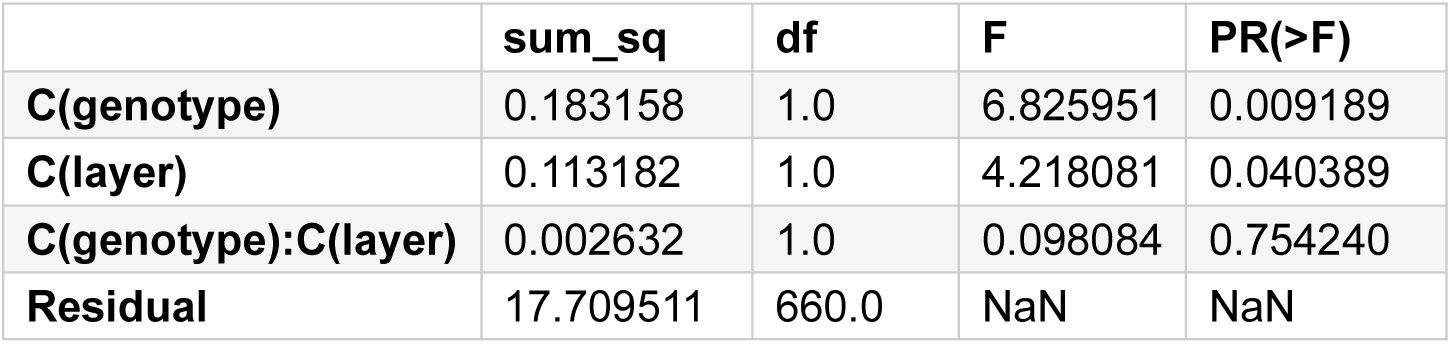
2 way ANOVA analysis of astrocyte contact fraction

**Table S2.** Glycogen granules counts in astrocytic processes

| Layer | Genotype | False<br>(no dark granules) | True<br>(dark granules) | All |
| --- | --- | --- | --- | --- |
| Layer II | KO | 50 | 112 | 162 |
|  | WT | 176 | 18 | 194 |
| Layer V | KO | 57 | 109 | 166 |
|  | WT | 114 | 28 | 142 |
| All |  | 397 | 267 | 664 |

